# DISTINCT ACTIN-DEPENDENT NANOSCALE ASSEMBLIES UNDERLIE THE DYNAMIC AND HIERARCHICAL ORGANIZATION OF E-CADHERIN

**DOI:** 10.1101/851899

**Authors:** Rumamol Chandran, Girish Kale, Jean-Marc Philippe, Thomas Lecuit, Satyajit Mayor

## Abstract

Intercellular adhesion mediated by E-cadherin is pivotal in maintaining epithelial tissue integrity and for tissue morphogenesis. Adhesion requires homophilic interactions between extracellular domains of E-cadherin molecules from neighboring cells. The interaction of its cytoplasmic domains with the cortical acto-myosin network, appears to strengthen adhesion, although, it is unclear how cortical actin affects the organization and function of E-cadherin dynamically. Here we use the ectopic expression of *Drosophila* E-cadherin (E-cad) in larval hemocytes, which lack endogenous E-cad, to recapitulate functional cell-cell junctions in a convenient model system. We used fluorescence emission anisotropy-based microscopy and Fluorescence Correlation Spectroscopy (FCS) to probe the nanoscale organization of E-cad. We find that E-cad at cell-cell junctions in hemocytes exhibits a clustered *trans-*paired organization, similar to that reported for the adherens junction in the developing embryonic epithelial tissue. Further, we find that extra-junctional E-cad is also organized as relatively immobile nanoclusters as well as diffusive and more loosely packed oligomers and monomers. These oligomers are promoted by *cis-*interactions of the ectodomain and, strikingly, their growth is constantly counteracted by cortical actomyosin. Oligomers in turn assist in generating nanoclusters that are stabilized by cortical acto-myosin. Thus, actin remodels oligomers and stabilizes nanoclusters, revealing a requirement for actin in the dynamic organization of E-cad at the nanoscale. This dynamic organization is also present at cell-cell contacts (junction), and its disruption affects junctional integrity in the hemocyte system, as well as in the embryo. Our observations uncover a hierarchical mechanism for the nanoscale organization of E-cad, which is necessary for dynamic adhesion and maintaining junctional integrity in the face of extensive remodeling.

## INTRODUCTION

Cadherins are transmembrane proteins required for the formation of multicellular tissues in organisms [1–4]. Classical cadherins contain amino-terminal extracellular domain or ectodomains followed by transmembrane domain and an intracellular domain. Typically, ectodomains of Cadherins engage in homophilic interactions with proteins present on adjacent cells (*trans* interactions), or the membrane of the same cell (*cis* interactions).

Work from a number of laboratories have indicated that intracellular interactions of the cytoplasmic domains of the E-cadherin with cortical acto-myosin network are a key to modulated cell-cell adhesion [5, 6]. The cytoplasmic tail of E-cadherin is known to interact with Catenin family proteins, α- and β-catenin [7–9]. E-cadherins bind directly to β-catenin which acts as an adaptor protein for the F-actin-binding α-catenin, thereby linking up to the cortical acto-myosin network [10–13]. α-catenin is also reported to have interactions with other F-actin binding proteins such as vinculin, EPLIN, and α-actinin thereby providing a number of linkages of the Cadherin-Catenin complex to cortical actin [14, 15].

Cell studies also show that the E-cadherin adhesion complex is under constitutive acto-myosin generated tension [6, 16]. E-cadherin and its associated components are also necessary for the integrity of the epithelial tissue in embryos of invertebrates such as *Drosophila* and vertebrates. The removal of adhesion complex components results in collapse of the tissue and in severe developmental defects [2,4,17]. This poses limitation for using the embryo system in exploring the mechanism behind the organization and function of the E-cadherin adhesion complex in a physiological context.

Studies from a number of groups have provided evidence for oligomers of E-cadherin at the cell membrane and its regulation by endocytosis, which in turn regulates the levels of E-Cadherin and the dynamics of Cadherin adhesion [18–21]. Super-resolution microscopy studies have also revealed the existence nanoscale clustering of E-cadherin at the cell-cell junctions directed by the actin mesh [22]. Recent studies using iPALM 3D-super-resolution microscopy characterized the nanoscale architecture of cadherin adhesions where they mapped the physical organization of *trans*-homophilic E-cadherin clusters [23, 24].

Imaging studies of *Drosophila* E-cadherin (E-cad) tagged to GFP (E-cad::GFP) expressed in *Drosophila* epithelial tissue shows a typical pattern outlining cells [11]; see **Figure 1A, top panel**). Studies in *Drosophila* embryo have also shown that E-cad is organized as finite size clusters referred to as spot adherens junctions (SAJs) in scanning electron micrographs or E-cad clusters using confocal imaging (see cartoon in **Figure. 1B**). These E-cad clusters [24] co-localized with actin patches, where it exchanges with the E-cad pool surrounding the clusters (extra-junctional pool) [11]. There is also evidence for the *cis*-clustering of E-cadherin outside the cell-cell adhesion sites which regulates cell-shape dynamics in *C. elegans* zygote [25]. Recent studies focusing on mechano-transducing property of E-cadherin revealed that E-cadherin is under constitutive actomyosin generated tension both outside and at cell-cell junctions [26]. The junctional tension is increased when cell-cell contacts are under strain, suggesting the role of cadherins as mechano-sensors.

**Figure 1:**
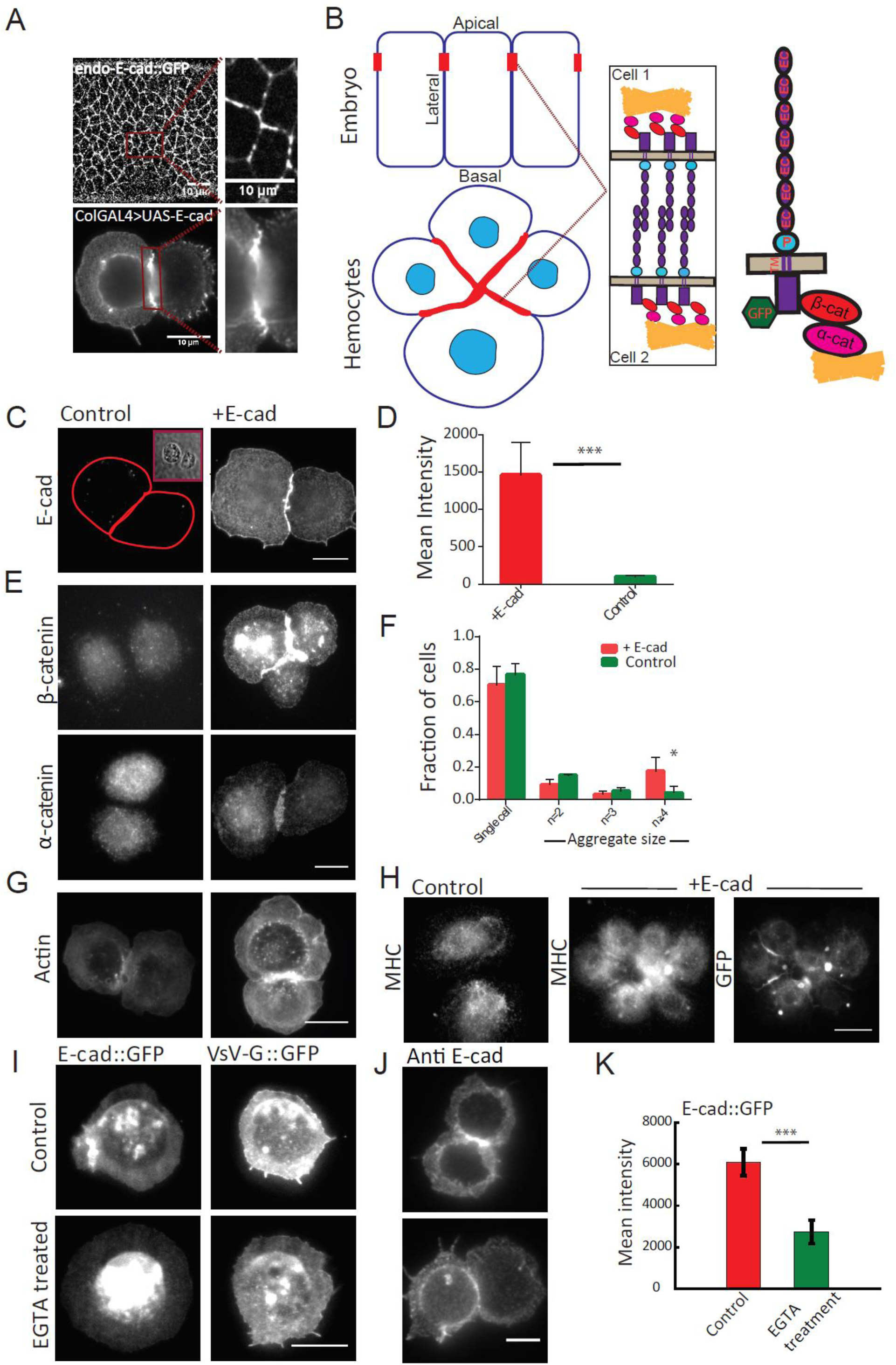
Functionally active E-cad on larval hemocytes membrane. (A) Images of E-cad::GFP expressing epithelial cells in an early developing embryo (left top) and membrane E-cadherin labeled using DCAD2 in hemocytes from wandering 3^rd^ instar larvae (left bottom); insets show magnified view of E-cad in embryos (right top) and hemocytes (right bottom). (B) Schematic showing *Drosophila* embryo epithelial cells and larval hemocytes forming cell-cell junctions with a zoomed view of the interactions happening at the junctions. Schematic showing *Drosophila* E-cadherin and its interaction machinery. (C) Distribution of E-cadherin in ectopic E-cad expressing and control hemocytes, as labeled with DCAD2 antibody with bright field image on the top right for the control cells. Scale bars, 10μm. (D) Histogram showing quantification of membrane pool of E-cad in the control and ectopic E-cad expressing hemocytes. (E) Immunostaining images of β-catenin and α-catenin in control and ectopic E-cad expressing hemocytes. Scale bars, 10μm. (F) Histogram showing the percentage of cells as single cells and cell aggregates in the control and ectopic E-cad expressing hemocytes. Data set contains normalized cell number from 5 control larvae and 6 ectopic E-cad expressing larvae and the bar represents the mean and the standard deviation in each condition. (G) Localization of actin in ectopic E-cad expressing and control hemocytes as labeled with Alexa-568 labeled phalloidin. Scale bars, 10μm. (H) 100X images of ectopic E-cad expressing and control hemocytes showing the localization of MHC using zipper antibody. Right most is the GFP channel of ectopic E-cad expressing hemocytes. (I) Representative TIRF images of untreated and EGTA treated cells in E-cad and VsV-G expressing hemocytes. Scale bars, 10μm. (J) Immunostaining images of membrane E-cad using DCAD2 antibody in ectopic E-cad expressing hemocytes in control and EGTA conditions. Scale bars, 10μm. (K) Histogram showing quantification of the membrane pool of E-cad using DCAD2 antibody in the control and EGTA treated cells in ectopic E-cad expressing hemocytes. The error bars depict standard deviation in (D) and (F) and standard error of the mean in (K). p values are calculated using Mann-Whitney U test. ns, p>0.05; *, p<0.05; **, p<0.01; ***, p<0.001

Here we have used the *Drosophila* larval hemocytes as a convenient model cell system [27], to address the role of molecular components in the establishment of a functional organization of E-cad. In this system we are able to explore the molecular and genetic basis for the organization of ectopically expressed E-cad. Circulating hemocytes from *Drosophila* 3^rd^ instar larvae expressing ectopic E-cad (using UAS-Gal4 system) form cell-cell junctions reminiscent of E-cadherin junctions in the embryo. This provides an optically and genetically tractable system to study the organization of E-cad in primary cells from various genetic backgrounds, expressing wild-type or mutant constructs of E-cad. The lamellar part of the hemocyte provides an ideal surface to study the organization of E-cad at extra-junctional regions, providing insights into the organization of E-cad at the relatively inaccessible apical surface of the embryonic epithelium.

Here we report that E-cad is hierarchically organized at multiple scales in the membrane of hemocytes, as well as in the junctional area in embryonic epithelial cells. Using fluorescence emission anisotropy-based microscopy we find that E-cad is *trans*-paired at the cell-cell junction, where it is localized in high density patches. Using fluorescence emission anisotropy based homo-FRET microscopy [28] we find that E-cad is organized as nanoscale clusters at the lamellar region of the hemocyte, and at extra-junctional apical membrane of the epithelial tissue in the *Drosophila* embryo. These nanoclusters require the α and β-Catenin complex and are also found at embryonic junctions. In addition, the nanoclusters are dependent on acto-myosin machinery for their formation, as well as ectodomain driven self-association of E-cad.

Fluorescence Correlation Spectroscopy (FCS) analysis of diffusive species in the membrane of cells [29, 30], show that E-cad forms diffusive oligomers due to the self-association property of its ectodomain, even in the absence of interactions with cytoplasmic acto-myosin. Finally, we find that extracellular domain association followed by nanocluster formation due to intracellular interaction with acto-myosin meshwork, is required for cell-cluster formation in circulating hemocytes suggesting that these processes strengthen the adhesion complex at the junctional membrane.

## RESULTS

### Functionally active E-cad on larval hemocytes membrane

E-cad is a junctional protein that may be visualized at the apical junction of embryonic epithelium by expression of E-cad::GFP at the endogenous locus [11]; **Figure 1A, top panel**). As is evident, the protein is present in concentrated large scale clusters at this surface as visualized from previous studies [24]. However, the topology of this tissue and the low level of endogenous expression of the protein at the apical membrane preclude a detailed quantitative analysis of its dynamic organization. To study the membrane organization of E-cad and its regulation by acto-myosin machinery we ectopically expressed E-cad in hemocytes derived from wandering 3^rd^ instar larvae, using UAS-GAL4 system as described in Experimental Procedures. Immunostaining with DCAD2 antibody revealed that the protein is undetectable in control cells, while present predominantly at the cell-cell junction of E-cad expressing cells **(Figure 1A, bottom panel, 1C, 1D)**. Further, immunostaining the adaptor proteins, α and β-catenin, showed their re-localization to the cell-cell junction in association with E-cad, in comparison to the cytoplasmic localization of these proteins in hemocytes derived from control larvae **(Figure 1E)**. Interestingly immunostaining of E-cad expressing cells showed increased expression of β-catenin compared to the wild type cells that could be due to the stabilization of the protein at the cell-cell junctions (compare images in **Figure 1E, top panels**). Immunostaining of actin **(Figure 1G)** and myosin heavy chain **(Figure 1H)** also showed the enrichment of acto-myosin at the cell-cell junctions of E-cad expressing cells compared to the control cells.

Ectopic expression of E-cad in hemocytes produces cell aggregates. A higher percentage of cells formed cell aggregates in E-cad expressing cells compared to the wild type cells consistent with the presence of functional E-cadherin on the membrane **(Figure 1F)**. The extracellular domains (EC repeats) of E-cad are calcium sensitive and binding of Ca^+2^ ions is necessary to rigidify ectodomains, which in turn helps in creating stable *trans* homophilic interactions at the junctions [31, 32]. Similar to E-cad in epithelial cells, E-cad expression at the hemocyte cell surface and its ability to create stable *trans* homophilic interactions at the junctions is dependent on Ca^+2^ **(Figure 1I-1K)**.

These observations provide evidence that functionally active E-cad is expressed at the plasma membrane of hemocytes. This protein interacts with and recruits the adaptor proteins, α- and β-catenin, as well as associates with the acto-myosin machinery (see cartoon in **Figure 1B**). The junctional localization of α- and β-catenin in hemocytes, in the presence of ectopic E-cad, provide a good model system to study the organization of E-cad at cell–cell adhesions.

### Emission anisotropy measurements reveal *trans*-pairing of E-cad at cell-cell junctions

We next studied the organization of fluorescently-tagged E-cad ([33]; endo-E-cad::GFP expressed at the endogenous locus) at the membrane of embryonic ectodermal cells. We imaged endo-E-cad::GFP, expressed in these epithelial cells during gastrulation movements of early embryonic development, using a spinning disc microscope equipped with the capacity to measure fluorescence intensity and emission anisotropy. In this method, using a customized confocal spinning disc microscope, polarized excitation is used to illuminate the sample and the resultant fluorescence emission is collected in parallel and perpendicular orientations (with respect to the axis of excitation polarization) to determine emission anisotropy. Rotational diffusion of the fluorophore will reduce emission anisotropy (EA), a situation also referred to below as depolarization of EA. Conversely restriction of this motion (for example by increasing the rotational mass of the protein complex) will result in an increase of emission anisotropy, also referred to as polarization of EA.

Endo-E-cad::GFP is mainly present in the apical region of the epithelium in two locations, at cell adherens junctions as well as in the free membrane at the apical surface. We observed visibly discernable spots of GFP tagged E-cad at cell junctions and a more uniform distribution in the apical membrane of the epithelium as previously described [11]. The spots represent *trans*-clusters of E-cad in-between cells, as observed previously [24].

We next measured the emission anisotropy of endo-E-cad::GFP in the embryonic tissue, at junctions and at the apical surface (as outlined in **Figure S1A)** using our customized confocal spinning disc microscope. Analysis of the anisotropy values of the endo-E-cad::GFP at the junctional punctae showed that it is measurably polarized compared to the endo-E-cad::GFP present at the free apical surface membrane **(Figure 2A-2B)**. As a control we examined the anisotropy values of ectopically expressed GFP-tagged transmembrane protein VsV-G. While this GFP-tagged protein did not form any visible punctae at the cell-cell junction, its anisotropy values in the apical surface and junctional plane of the epithelial cells are similar to each other **(Figure S1C, S1D)**.

**Figure 2:**
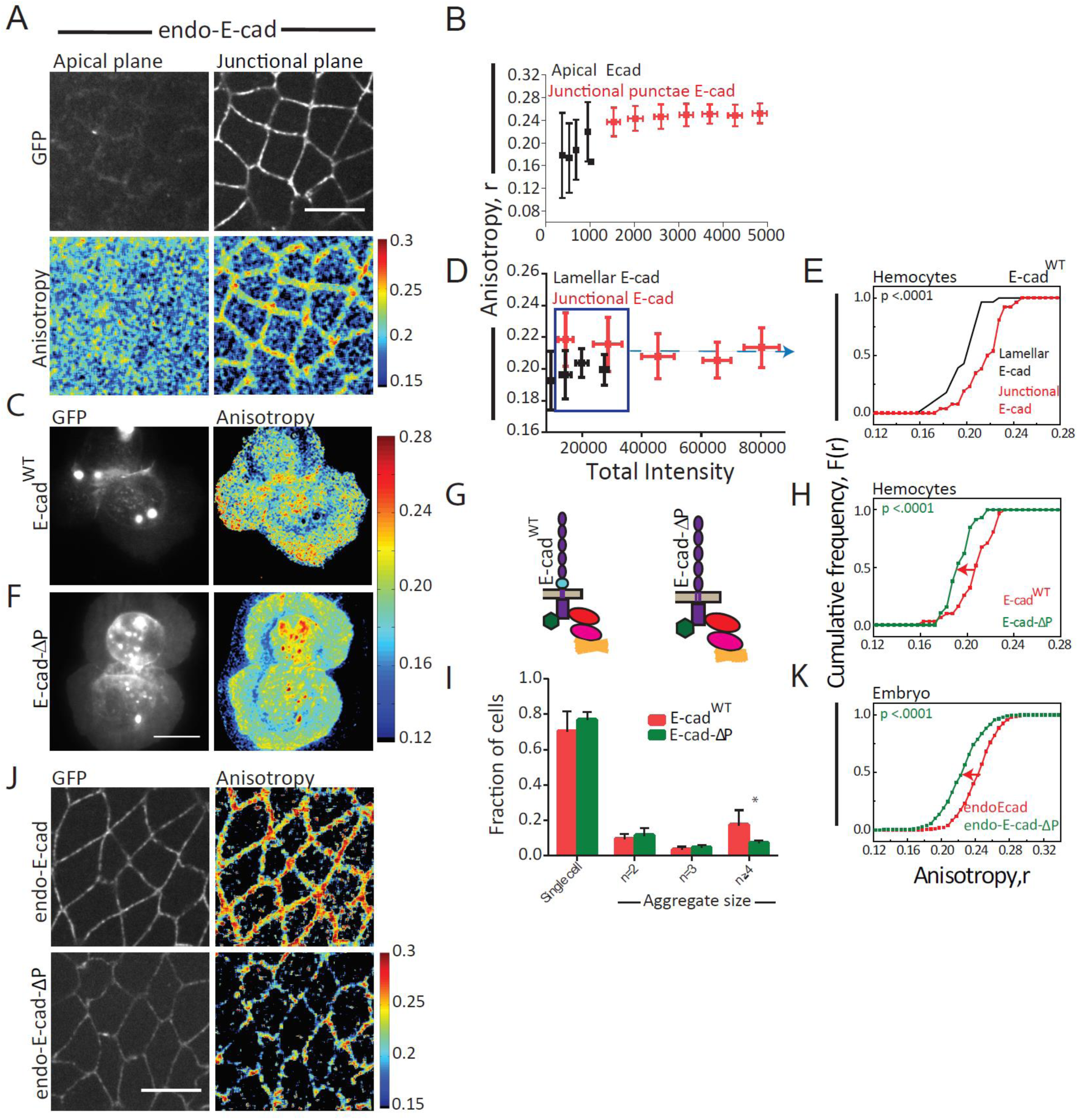
Nanoscale organization of E-cad in embryo and larval hemocytes probed by Homo-FRET microscopy. (A) Total intensity and anisotropy images of endo E-cad::GFP in the apical and junctional plane in the early gastrulation stage of the embryo. Scale bars, 10μm. (B) Intensity versus anisotropy plots of endo-E-cad in the apical and junctional plane from similar stage embryos (N=5). (C) Total intensity and anisotropy images of E-cad^WT^::GFP expressing hemocytes. Scale bars, 10μm. (D) Scatter plot of intensity versus anisotropy of lamellar and junctional E-cad^WT^. (E) Cumulative frequency distributions of anisotropy values shown in (D) are plotted for lamellar and junctional E-cad^WT^ of similar intensity range. (F) Total intensity and anisotropy images of E-cad-ΔP::GFP mutant expressing hemocytes Scale bars, 10μm. (G) Schematic of E-cad^WT^::GFP and E-cad-ΔP::GFP proteins. (H) Cumulative frequency distributions of junctional anisotropy values of E-cad^WT^ and E-cad-ΔP expressing hemocytes. (I) Histogram showing the percentage of cells as single cells and cell aggregates in E-cad^WT^::GFP and E-cad-ΔP::GFP expressing hemocytes. Data set contains normalized cell number from 6 E-cad^WT^ larvae and 4 E-cad-ΔP larvae and the bar represents the mean and the standard deviation in each condition. (J) Total intensity and anisotropy images of endo-E-cad::GFP and endo-E-cad-ΔP::GFP expressing embryos. Scale bars, 10μm. (K) Cumulative frequency distributions of anisotropy values of junctional E-cad in endo-E-cad (N=5) and endo-E-cad-ΔP (N=5) expressing embryos. p values are calculated using Mann-Whitney U test. ns, p>0.05; *, p<0.05; **, p<0.01; ***, p<0.001

Perturbations that may result in defects in junctional stability in the embryo, limits our ability to study different E-cad mutant isoforms ectopically expressed in a background that lacks endogenous E-cad. Therefore, we further characterized the organization of E-cadherin and its variants in *Drosophila* larval hemocytes. We imaged GFP-tagged E-cad [34] in hemocytes using EA based TIRF microscopy [EA-TIRFM]; [35]. We observed that the fluorescence emission anisotropy of E-cad^WT^ in lamella (corresponding to the apical surface in the embryo) is also depolarized compared to its junctional counterparts, across a similar total intensity range of membrane localized E-cad **(Figure 2C-2E)**. We also observed that the lamellar E-cad^WT^ emission anisotropy is similar in isolated cells as well as in the clustered cells **(Figure S1E).**

As mentioned above, extracellular *trans*-interactions of the junctional protein in opposing membranes could restrict rotational movement of the proteins resulting in more polarized emission of the ensemble of fluorescent molecules at the junction. Since the fluorescence emission anisotropy of an ensemble of fluorophores is a linear combination of the emission anisotropy of individual fluorophores, this led us to test whether the higher emission anisotropy values of the protein at the junctions is due to *trans*-pairing with E-cad in opposing membranes. For this purpose, we utilized a mutant of E-cad that is expected to have defects in homophilic interactions [36]. This mutant has deletions in the proximal domain of E-cad, which comprises of NC domain, CE domain and LG domain. Such a mutation affects tissue integrity in cells with high actomyosin contractility such as in the *Drosophila* mesoderm [36]. When the proximal domain deleted E-cad tagged to GFP (schematic in **Figure 2G**; E-cad-ΔP::GFP, [36]) was expressed in hemocytes, we found that the E-cad-ΔP expressing hemocytes exhibited fewer cell aggregates as compared to those formed by E-cad^WT^ **(Figure 2I)** at similar or higher levels of protein expression, consistent with its defective capacity to form homophilic interactions. Importantly, emission anisotropy values of E-cad-ΔP at the junction were not as polarized compared to junctional E-cad^WT^ **(Figure 2F, 2H)**. When endo-E-cad-ΔP::GFP was expressed in the embryo in the background of untagged endogenous E-cad, the emission anisotropy of junctional endo-E-cad-ΔP was also much reduced compared to the endo E-cad::GFP expressed under same conditions (**Figure 2J, 2K)**. This suggests that the enhancement of emission anisotropy values of the junctional E-cad compared to that measured on free apical surface of the epithelial cells or the lamella of the hemocyte is due to the restriction in rotational mobility of the proteins. This effect is likely due to *trans*-pairing of at least a subset of E-cad proteins at the cell-cell junctions.

### E-cad is clustered at the nanoscale at the junction and in extra-junctional regions

To assess whether E-cad is tightly clustered at the junction at the nanoscale as suggested [22, 24], we utilized the propensity of fluorescence emission anisotropy to be reduced by FRET between like species (homo-FRET). The extent of reduction in this value of emission anisotropy is a measure of FRET efficiency between an ensemble of fluorescent molecules [37]. Construction of emission anisotropy maps of the fluorescent protein therefore provides a spatial distribution of regions of the cell membrane where the protein is organized at different scales. Correspondingly, polarized (high) anisotropy values (depicted as red pixels in the emission anisotropy maps) reflect less *cis*-clustered or a more rotationally constrained fluorophore (eg. by *trans*-interactions) and depolarized (low) anisotropy values (depicted as blue pixels in the emission anisotropy maps), also indicate the location of *cis*-clustered species **(Figure S1B)**.

Since the depolarization is dependent on the presence of neighboring fluorophores, the extent of homo-FRET may be assessed by a photobleaching assay, wherein photobleaching will result in a steady increase in anisotropy values upto the value of the monomeric species (see schematic in **Figure 3A**). However, if proteins which are dispersed far apart as monomers on the membrane and/or are only rotationally constrained (e.g. by *trans-*pairing), they will not exhibit a change in anisotropy values upon photobleaching. To assess the extent of homo-FRET, we photobleached for E-cad^WT^::YFP **(Figure 3B)** expressed in hemocytes, and monitored emission anisotropy values of lamellar and junctional E-cad. In this assay, as outlined by the schematic **(Figure 3A; see also** [38]), photo-bleaching of E-cad^WT^::YFP in the lamellar and the junctions **(Figure 3C, 3D)** both exhibited a linear increase in anisotropy values, consistent with a fraction of the E-cad^WT^::YFP species engaging in homo-FRET at these locations. The linear nature of the increase in anisotropy upon photobleaching for E-cad^WT^::YFP at the lamella and junction is characteristic of the presence of a fraction of dispersed nanoclusters of E-cad^WT^::YFP, and not a uniform large scale patch of densely packed molecules [38, 39]. These data are entirely consistent with the observation of a collection of nanoscale clusters as evidenced from STORM studies in E-cad expressed in mammalian epithelia [22] and from PALM in *Drosophila* embryos [24], of the junctional protein. Since the YFP moiety is located in the cytoplasmic domain of the proteins, this implies that E-cad are close enough to undergo homo-FRET [38], and therefore engage in measurable *cis*-interactions in the junction as well as in the lamella [25].

**Figure 3:**
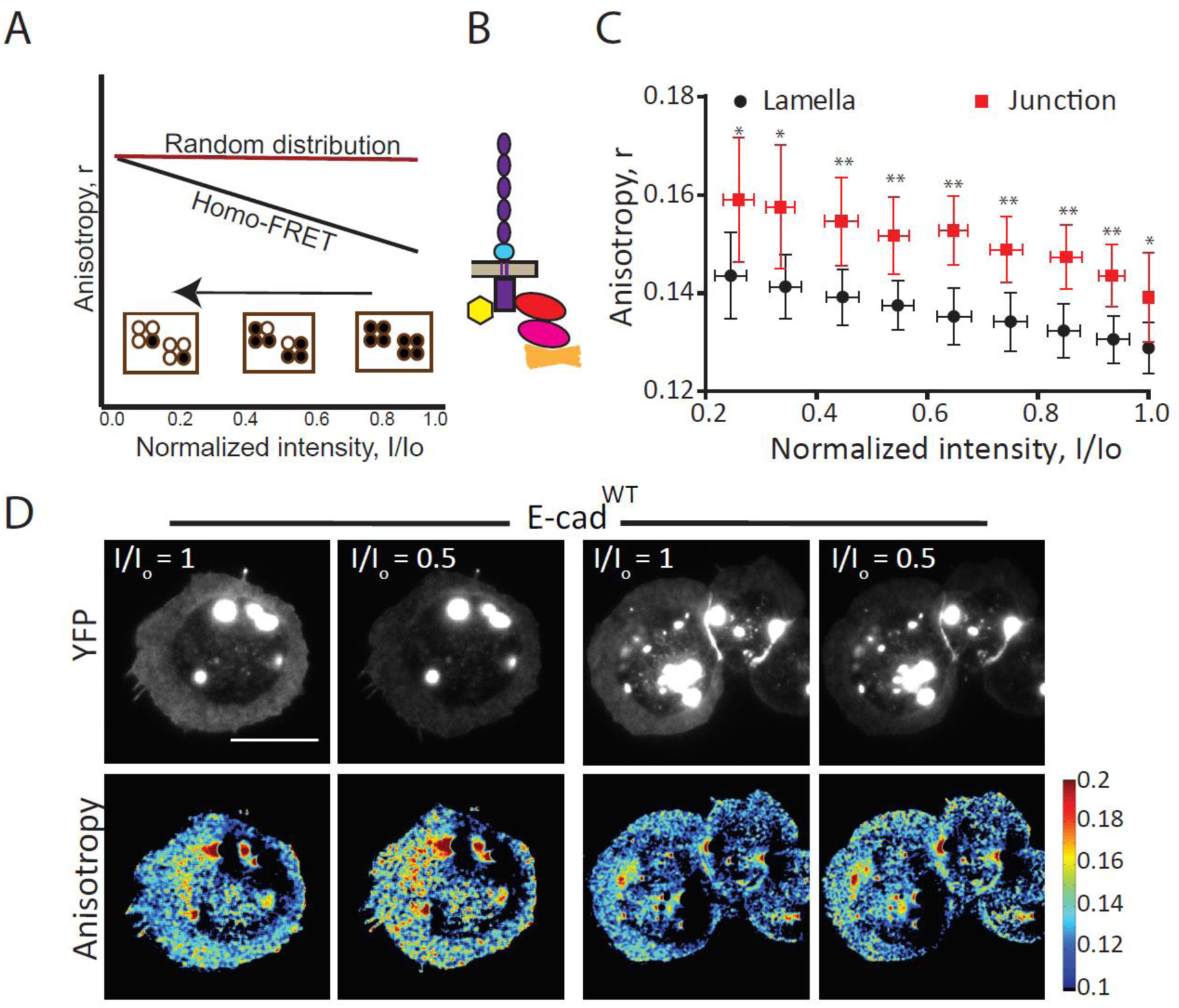
Nanoscale clustering of junctional and extra-junctional E-cad in hemocytes. (A) Schematic of the photobleaching assay is shown wherein the filled circles represent the molecules which can fluoresce and the open circles represent the bleached fluorophore. A typical photobleaching profile of nanoclustered molecules is shown in black line and the red line represents the scenario where there are less nanoclusters. The anisotropy values of ROIs are plotted against the intensity values which are normalized to the starting total intensity value of ROI. (B) Schematic of E-cad^WT^::YFP construct used for photobleaching experiments. (C) Intensity versus anisotropy plots of lamellar and junctional E-cad in E-cad^WT^ expressing cells depicting the changes in anisotropy values upon photobleaching. Each data point represents average anisotropy values taken from multiple regions from different cells with standard deviation for the corresponding intensity bins normalized to the starting intensity value. (D) Total intensity and anisotropy images of E-cad^WT^::YFP expressing cells at the beginning of photobleaching and at 50% photobleaching for the single cells (left panel) and cell cluster (right panel). Scale bars, 10μm. P values are calculated using Mann-Whitney U test. Ns, p>0.05; *, p<0.05; **, p<0.01; ***, p<0.001

Comparing the profile of the change in emission anisotropy of junctional and lamellar protein due to photobleaching, it is evident that they follow parallel paths, while the junctional anisotropy is always higher in value than the lamellar anisotropy for the entire photobleaching process (**Figure 3C; compare lamella and junction)**. This is consistent with the more restricted rotational mobility of the protein engaged in *trans-*pairing interactions at the junctions compared to that at the free surface of the lamella. While the existence of very close packed interactions of the *trans-*paired E-cad at the junctions was expected, the detection of homo-FRET between E-cad molecules in the lamellar region is surprising. Recent studies on adhesion-independent cadherin clusters in the *C. elegans* embryo [25] have also provided evidence for the existence of extra-junctional clusters of E-cadherin.

### Adhesion complex is required for nanoscale *cis*-clusters of E-cad at the lamella

The hemocyte system we have developed here provides a good opportunity to explore the molecular mechanism of the nanoclustering of extra-junctional lamellar E-cad. *Cis-* clustering of mammalian E-cad at extra-junctional sites has been reported to be mediated mainly by the cytoplasmic domain, with some contribution from homophilic interactions of the extracellular domain in increasing the packing density of the protein [22]. We first addressed if this type of clustering requires the recruitment of the adhesion complex machinery by utilizing an E-cad mutant defective in binding β-catenin (see schematic in **Figure 4A)**. E-cad binds β-catenin directly and recruits the actin binding protein, α-catenin, to the complex. An E-cad mutant that lacks β-catenin binding domain (E-cad-Δβ; [40]) was expressed in hemocytes, to assess if the interaction with the adaptor proteins is necessary for clustering of lamellar E-cad. Immunostaining of the E-cad-Δβ expressed in hemocytes showed that it does not recruit β-catenin to the cell-cell junctions, compared to the junctional recruitment of β-catenin in the presence of wild type protein, confirming the loss of interaction of the mutant protein with β-catenin **(Figure S1F)**.

**Figure 4:**
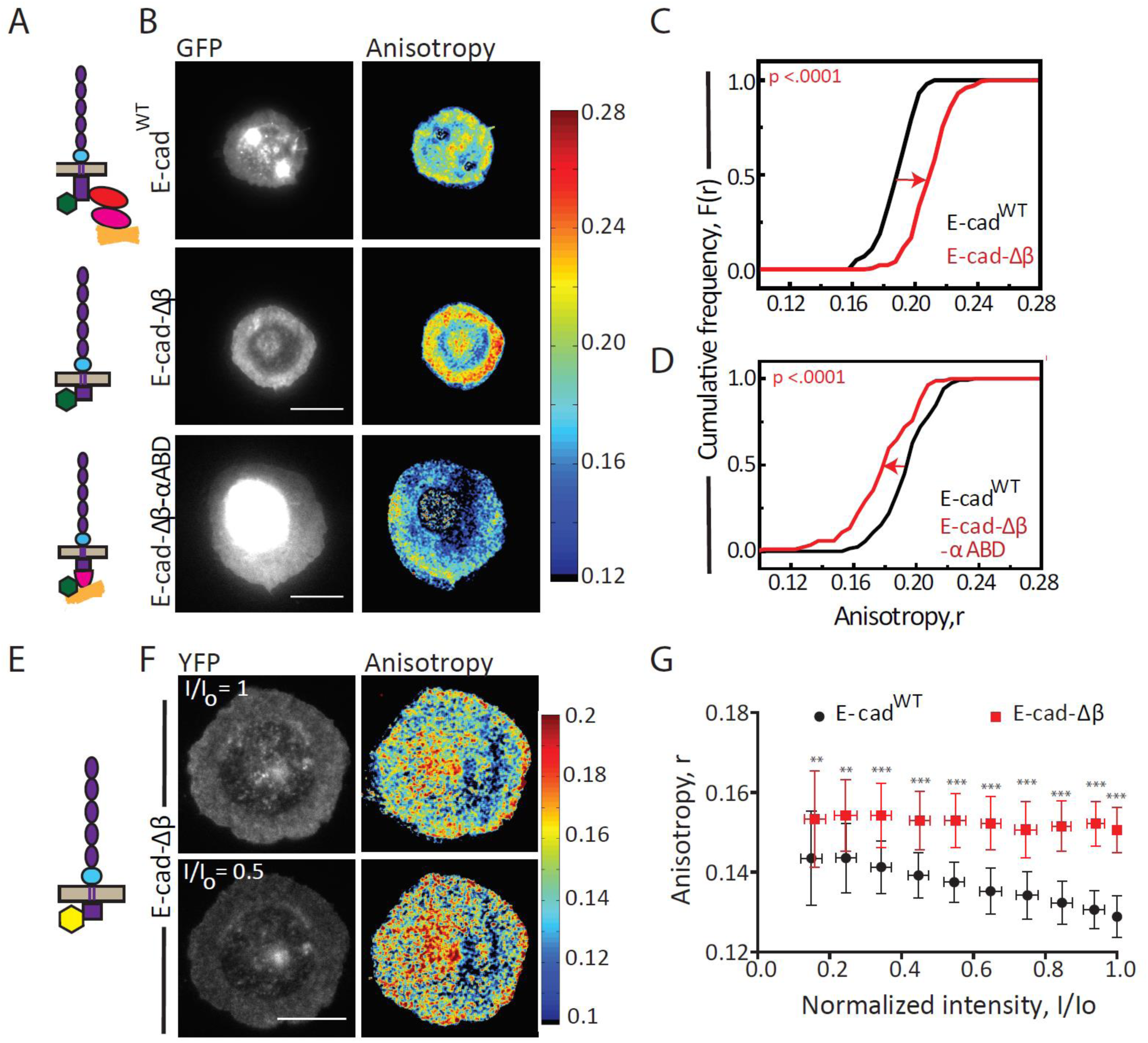
Adaptor complex regulates lamellar E-cad organization. (A) Schematic of E-cad mutant constructs used in the study. (B) Total intensity and anisotropy images of E-cad^WT^::GFP (top), E-cad-Δβ::GFP (middle) and E-cad-Δβ-αABD::GFP (bottom) expressing hemocytes. Scale bars, 10μm. (C) Cumulative frequency distributions of lamellar anisotropy values of E-cad^WT^ compared to E-cad-Δβ and (D) E-cad^WT^ compared to E-cad-Δβ-αABD expressing hemocytes. (E) Schematic of E-cad-Δβ::YFP construct used for photobleaching experiments. (F) Total intensity and anisotropy images of E-cad-Δβ::YFP expressing cells at the beginning of photobleaching and at 50% photobleaching. Scale bars, 10μm. (G) Intensity versus anisotropy plots of E-cad::YFP in E-cad^WT^ and E-cad-Δβ expressing cells depicting the changes in anisotropy values upon photobleaching. P values are calculated using Mann-Whitney U test. Ns, p>0.05; *, p<0.05; **, p<0.01; ***, p<0.001

Lamellar E-cad-Δβ::GFP has emission anisotropy that is significantly polarized compared to E-cad^WT^ protein **(Figure 4B middle panel, 4C)**, and importantly lamellar E-cad-Δβ::YFP (see schematic in **Figure 4E)** did not show a detectable change in emission anisotropy values upon photobleaching (**Figure 4F, 4G**) consistent with the lack of nanoclustering of this mutant protein on the lamella membrane. These data suggest that cytoplasmic interaction with β-catenin is necessary for the nanoscale lamellar E-cad organization.

To address whether interaction with α-catenin facilitates this nanoscale organization, we expressed chimeric proteins of E-cad-Δβ with full length or truncated α-catenin domains that carries only the actin binding domain of α-catenin in hemocytes [domain structure of *Drosophila* α-catenin schematized in **Figure S1G;** [40, 41]]. Only E-cad-Δβ with α-catenin actin binding domain (E-cad-Δβ-αABD) is trafficked to the cell surface in hemocytes, whereas the other E-cad-Δβ chimeric proteins including E-cad-Δβ with full length α-catenin are not delivered to the cell surface. They are instead sequestered in intracellular compartments **(Figure S1H**). This allowed us to only compare the properties of E-cad-Δβ and E-cad-Δβ-αABD with E-cad^WT^.

Lamellar E-cad-Δβ-αABD::GFP emission values were marginally depolarized compared to E-cad^WT^ **(Figure 4B bottom panel, 4D)**, and consequently significantly depolarized with respect to E-cad-Δβ, suggesting that stable association with actin binding domain of α-catenin is required for the clustering of E-cad on the lamella membrane. Together these results suggest that at least a fraction of lamellar E-cad exists as nanoclusters and its ability to interact with the adaptor proteins, β- and α-catenin, likely drives the nanoclustering of E-cad on the lamellar membrane.

### Acto-myosin activity organizes lamellar E-cadherin into nanoclusters

E-cad is known to associate with acto-myosin machinery via its interaction with α-catenin [10, 42], therefore we next probed if perturbations of the acto-myosin machinery could alter the nanoclustering characteristics of lamellar E-cad^WT^. Disruption of actin machinery by actin depolymerizing drug, Latrunculin A showed higher anisotropy values compared to control cells **(Figure 5A left panel, 5B)**, and genetic perturbations of non-muscle myosin 2, myosin regulatory light chain (MRLC) and myosin heavy chain (MHC), results in loss of nanoclustering of lamellar E-cad **(Figure 5A right panel, 5C)**. Pre-treatment of the cells with ROCK inhibitor, H1115 [43] also resulted in an increase in the anisotropy values, which was consistent with loss of nanoclusters of lamellar E-cad **(Figure 5D, 5E)**. We conclude that the acto-myosin network and the regulator ROCK are all part of the machinery that mediates nanoclustering of lamellar E-cadherin.

**Figure 5:**
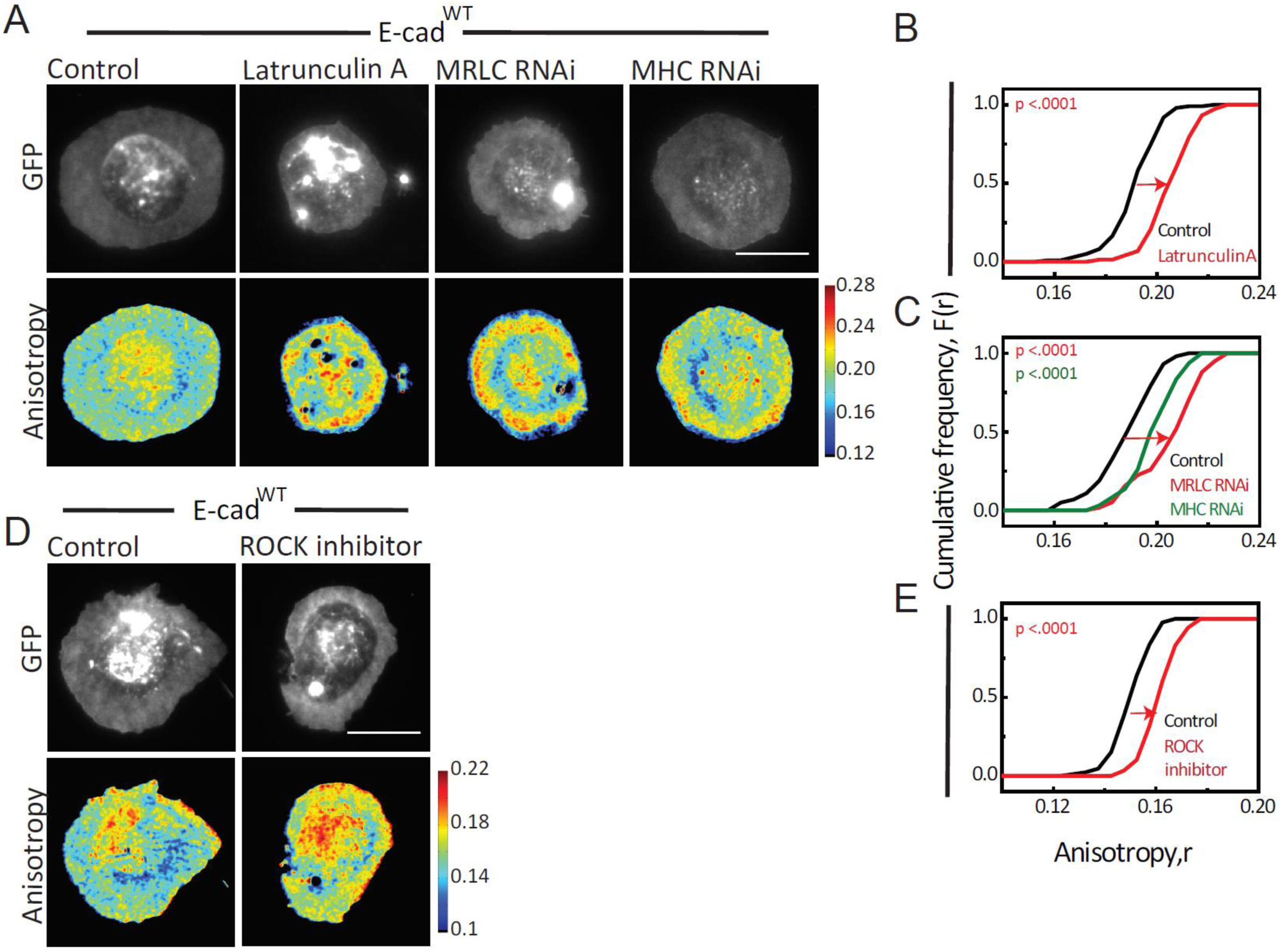
Acto-myosin activity organizes lamellar E-cad. (A) Total intensity and anisotropy images of E-cad^WT^::GFP in control, Latrunculin A treated, MRLC RNAi and MHC RNAi expressing cells. Scale bars, 10μm. (B) Cumulative frequency distributions of anisotropy values of lamellar E-cad^WT^ in control and Latrunculin A treated cells. (C) Cumulative frequency distributions of anisotropy values of lamellar E-cad^WT^ in control, MRLC RNAi and MHC RNAi expressing hemocytes. (D) Total intensity and anisotropy images of E-cad^WT^::GFP in control and ROCK inhibitor treated hemocytes. Scale bars, 10μm. (E) Cumulative frequency distributions of anisotropy values of E-cad^WT^ in control and ROCK inhibitor treated hemocytes. P values are calculated using Mann-Whitney U test.

### Inside out interaction is essential for cell clustering and junctional organization

We next investigated whether the machinery regulating nanoclustering of lamellar E-cad also mediates the organization of junctional protein by investigating the organization of E-cad-Δβ. As established above, the anisotropy value of junctional E-cad-Δβ is more polarized than E-cad^WT^ at junctions, indicating a reduction in nanoclustering of E-cad-Δβ protein compared to E-cad^WT^ **(Figure 6A, 6B)**. By contrast, unlike the lamellar E-cad-Δβ, we see an increase in anisotropy of E-cad-Δβ::YFP upon photobleaching indicative of the presence of dense packed E-cad-Δβ species capable of homo-FRET **(Figure 6H red plot)**. The profile of photo-bleaching is, however, distinct from that obtained with E-cad^WT^::YFP (**Figure 6H black plot)**. The lack of a change in anisotropy until significant fraction of the fluorophore is bleached (30%), followed by a transition to a rise in anisotropy is entirely consistent with that expected from dense patches or aggregates much larger than the range of the homo-FRET process [Forster’s radius∼ 6 nm; [38]]. Thus, while nanoclustering is altered at the junctions when the cytoplasmic connection is perturbed, we suggest that *trans-*interactions are sufficient to promote a dense association of the E-cad proteins in larger aggregates.

**Figure 6:**
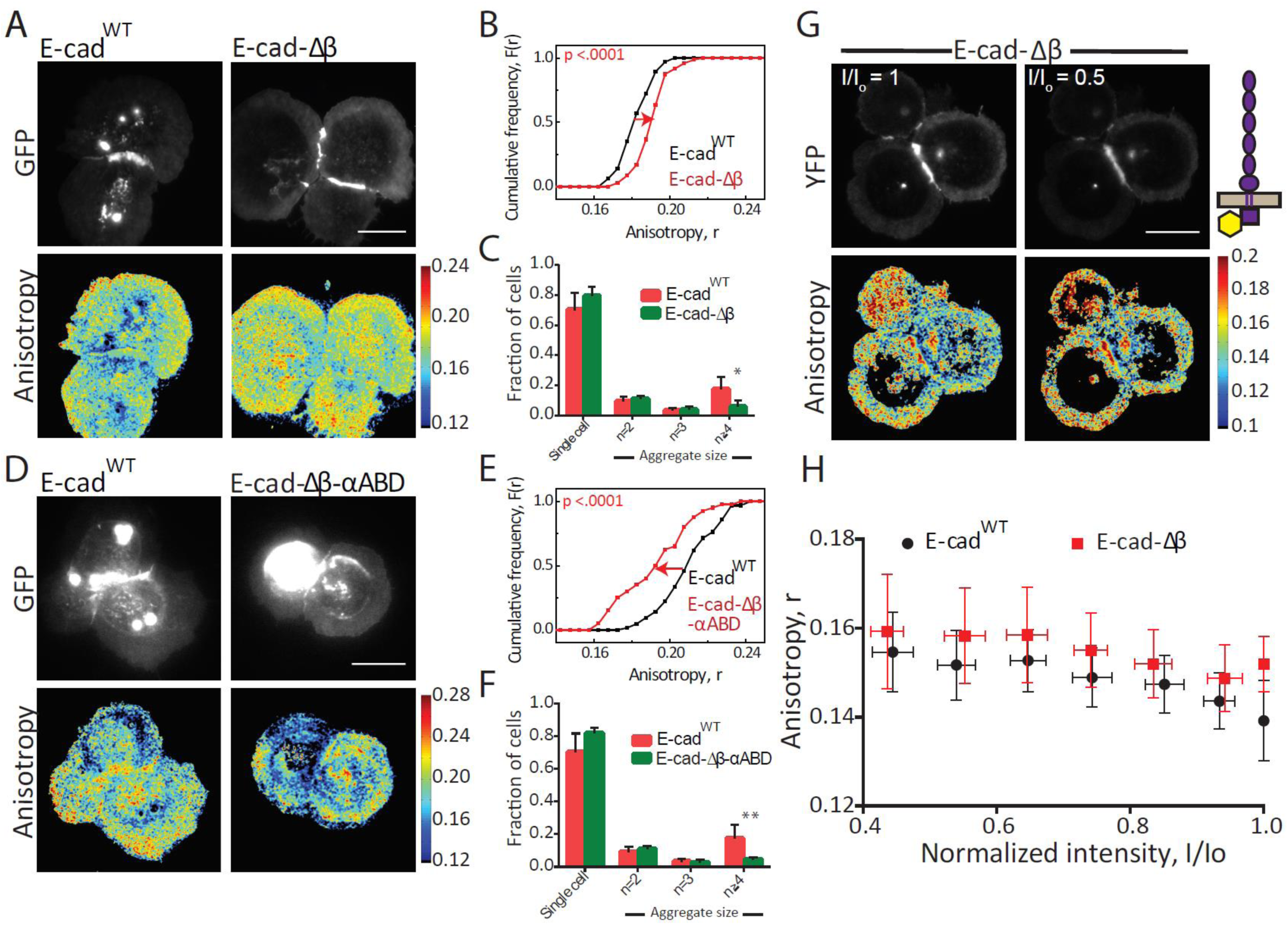
Adaptor protein dependent clustering of junctional E-cad. (A) Total intensity and anisotropy images of E-cad^WT^::GFP and E-cad-Δβ::GFP expressing cells. Scale bars, 10μm. (B) Cumulative frequency distributions of anisotropy values of junctional E-cad in E-cad^WT^ and E-cad-Δβ expressing hemocytes. (C) Histogram showing percentage of cell aggregates in E-cad^WT^::GFP and E-cad-Δβ::GFP expressing cells. Data set contains normalized cell number from 6 E-cad^WT^ larvae and 7 E-cad-Δβ larvae and the bar represents the mean and the standard deviation in each condition. (D) Total intensity and anisotropy images of E-cad^WT^::GFP and E-cad-Δβ-αABD::GFP expressing hemocytes. Scale bars, 10μm. (E) Cumulative frequency distributions of anisotropy values of E-cad in E-cad^WT^ and E-cad-Δβ-αABD expressing hemocytes. (F) Histogram showing percentage of cell aggregates in E-cad^WT^::GFP and E-cad-Δβ-αABD::GFP expressing cells. Data set contains normalized cell number from 6 E-cad^WT^ larvae and 5 E-cad-Δβ-αABD larvae and the bar represents the mean and the standard deviation in each condition. (G) Total intensity and anisotropy images of and E-cad-Δβ::YFP expressing cells at the beginning of photobleaching and at 50% photobleaching. Scale bars, 10μm. (H) Intensity versus anisotropy plots of junctional E-cad in E-cad^WT^ and E-cad-Δβ expressing cells depicting the changes in anisotropy values upon photobleaching. Each data point represents average anisotropy values taken from multiple regions of different junctions with standard deviation for the corresponding intensity bins normalized to the starting intensity value. P values are calculated using Mann-Whitney U test.

In contrast, expression of E-cad-Δβ-αABD::GFP resulted in the lowering of the anisotropy of the junctional protein compared to E-cad-Δβ and E-cad^WT^, indicating that the nanocluster organization of E-cad at the junctions is restored by linking it up to actin machinery, confirming the role of a dynamic acto-myosin machinery in the nanoscale organization of E-cad at the junction **(Figure 6D, 6E)**. Treatment with Latrunculin also increased emission anisotropy at the junction, consistent with a reduction of the nanoclustered organization of E-cad at the junction **(Figure S2A, S2B)**.

The role of acto-myosin in nanoclustering at the junction was addressed using chemical inhibitors and genetic perturbations. RNAi perturbations of non-muscle myosin 2 (MLC and MHC) showed polarized anisotropy values at the junctions **(Figure S2C, S2D)**. The inhibition of ROCK using the small molecule, H1152 also results in loss of clustering, confirming the acto-myosin dependence of junctional E-cadherin organization **Figure S2E, S2F)**.

As already shown, ectopic E-cad^WT^ expressing cells form more cell aggregates compared to hemocytes lacking E-cad^WT^ (control cells, **Figure 1F)**. The cell adhesion ability was affected when E-cad interaction with adaptor complex proteins was disturbed as evidenced by a reduction in the percentage of cells forming aggregates. Ecad-Δβ or Ecad-Δβ-αABD expressing hemocytes had fewer cells forming aggregates compared to the wild type protein (**Figure 6C, 6F**). This strongly suggested that the cell adhesion property of E-cad is dependent on its finely tuned association with its intracellular complex components.

These results led us to conclude that junctional as well as lamellar E-cad organization or clustering mainly depends on its interaction with acto-myosin machinery via adaptor proteins β-catenin and α-catenin. Polarization or increase in anisotropy values at the junction and lamella upon perturbation of associated machinery supported the existence of nanoscale acto-myosin based organization of junctional E-cad, necessary for inside out regulation of cell-cell adhesion.

### Nanoclustering of E-cad in embryonic epithelial cells is acto-myosin dependent

We have shown in the hemocyte system, the lamellar and junctional E-cad form nanoclusters which are dependent on acto-myosin machinery. We have also observed a similar nanoscale organization of E-cad in epithelial cells of *Drosophila* embryo **(Figure 2A-2B**) where junctional anisotropy is more polarized than at the apical surface. We next addressed whether the same molecular machinery, which regulates the nanoscale organization of E-cad in hemocytes, also operates in the embryo. The role of myosin-II in the remodeling of epithelial cell junctions in *Drosophila* embryo has been studied extensively [21,43,44]. Therefore, we used ROCK perturbations to remove the myosin pool associated with the junctional E-cad protein and examined its nanoclustering. We observed an increase in anisotropy values, indicative of loss of clustering of the junctional E-cad in ROCK inhibitor H1152 injected embryos compared to water injected embryos **(Figure S3A left panel, S3B).** We further assessed the role of actin in junctional E-cad organization using Latrunculin A injections. An increase in anisotropy values in Lat A injected embryos compared to DMSO injected embryos, confirmed the role of acto-myosin network in regulating the organization of junctional E-cadherin **(Figure S3A right panel, S3C)**.

Together these results strongly support the idea that E-cad is also present as nanoclusters at cell junctions in the embryonic epithelium and is dependent on the acto-myosin networks. In parallel, extracellular *trans-*interactions of the junctional protein results in restricted rotational tumbling of the protein and polarization of anisotropy values at the cell-cell junctions when compared to the contact free region of the apical membrane.

### E-cad makes dynamic diffusive oligomers on the free-membrane

The actomyosin-sensitive nanoclusters of E-cad described above are expected to exhibit turnover rates on the order of 0.1-1 sec^−1,^ as noted from experiments characterizing the dynamics of such types of nanoclustering [45]. Coupled with the expectation that these nanoclusters are relatively immobile this turnover rate is likely to result in a pool of immobile E-cad on the time scales accessible in FRAP experiments conducted in area (∼3μm diameter). Consistent with this expectation, FRAP experiments on E-cad^WT^ in the hemocyte lamella, indicated a small but significant immobile pool along with a large mobile fraction on the time scale of FRAP, namely few tens of seconds **(Figure S4A, S4B)**. Consistent with the immobile pool arising from the actomyosin interactions, this pool was significantly reduced, when the FRAP profiles of E-cad-Δβ were analyzed **(Figure S4A, S4B)**. These data are consistent with the role of E-cad cytoplasmic association with acto-myosin and nanoclustering, as a reason for its immobilization.

However, ∼ 70% of the E-cad species is mobile, and might form higher order structures [19, 20]. To characterize this mobile pool of E-cad species we used FCS [[29, 46]; see also Supplementary Methods], which provides the number and the average brightness of the of diffusing entities in the confocal volume. Further, ascertaining the brightness of single cytoplasmic GFP molecules **(Figure S5H)**, would also allow us to estimate the oligomeric state of the GFP-tagged diffusing species, comprised of E-cad^WT^::GFP or its variants. We first compared the brightness of cytoplasmic GFP to VSV-G-GFP, a membrane anchored protein that is known to form a trimer [47]. The measured brightness of cytoplasmic GFP and VSV-G::GFP indicated that although VSV-G::GFP is brighter than GFP (∼ 10 cpm versus 6.2) its “effective brightness per molecule in the trimer” is about 54% of cytoplasmic GFP **(Figure S5I)**, (lower than the expected brightness of the diffusing VSV-G::GFP trimer which should be ∼ 3x cytoplasmic GFP). Assuming a similar “effective brightness per molecule” for individual E-cad (GFP) molecules, we can conclude that diffusing oligomers of the E-cad^WT^ resemble an oligomeric species that is tetrameric, on an average, since they are approximately 1.25-fold brighter than VSVG-GFP.

We next analyzed the brightness of E-cad^WT^ after depolymerizing F-actin using Latrunculin A (LatA), a treatment which disrupts nano-clustering. Surprisingly, E-cad^WT^ in LatA treated cells made brighter, i.e. larger oligomeric species, as compared to DMSO treated E-cad^WT^ expressing cells **(Figure 7A).** This indicated that the cytoplasmic interactions with F-actin counteracts the tendency of E-cad oligomers to grow larger. We next compared the brightness of diffusive oligomers made by E-cad^WT^ with that of E-cad-Δβ, where cytoplasmic interaction capacity with actin is also lost. These results resembled those obtained in LatA treated cells; the E-cad-Δβ diffusive species are brighter than the E-cad^WT^ species **(Figure 7B)**. To follow-up this result further, we also extended this analysis to E-cad-Δβ-αABD. The oligomeric state of E-cad-Δβ-αABD was similar to that of E-cad^WT^ **(Figure 7B)**. If indeed the diffusible oligomers were due to *cis-* interactions of the ectodomain, we hypothesized that the E-cad-ΔP would exhibit smaller oligomers, if any, since the extracellular interactions of E-cad-ΔP are likely to be weak. Consistent with our prediction, the oligomers of E-cad-ΔP were even smaller than E-cad^WT^ **(Figure 7B)**. This indicated an antagonism between extracellular and intracellular cytoplasmic connections to regulate the dynamic oligomerization of the mobile pool of E-cad.

**Figure 7:**
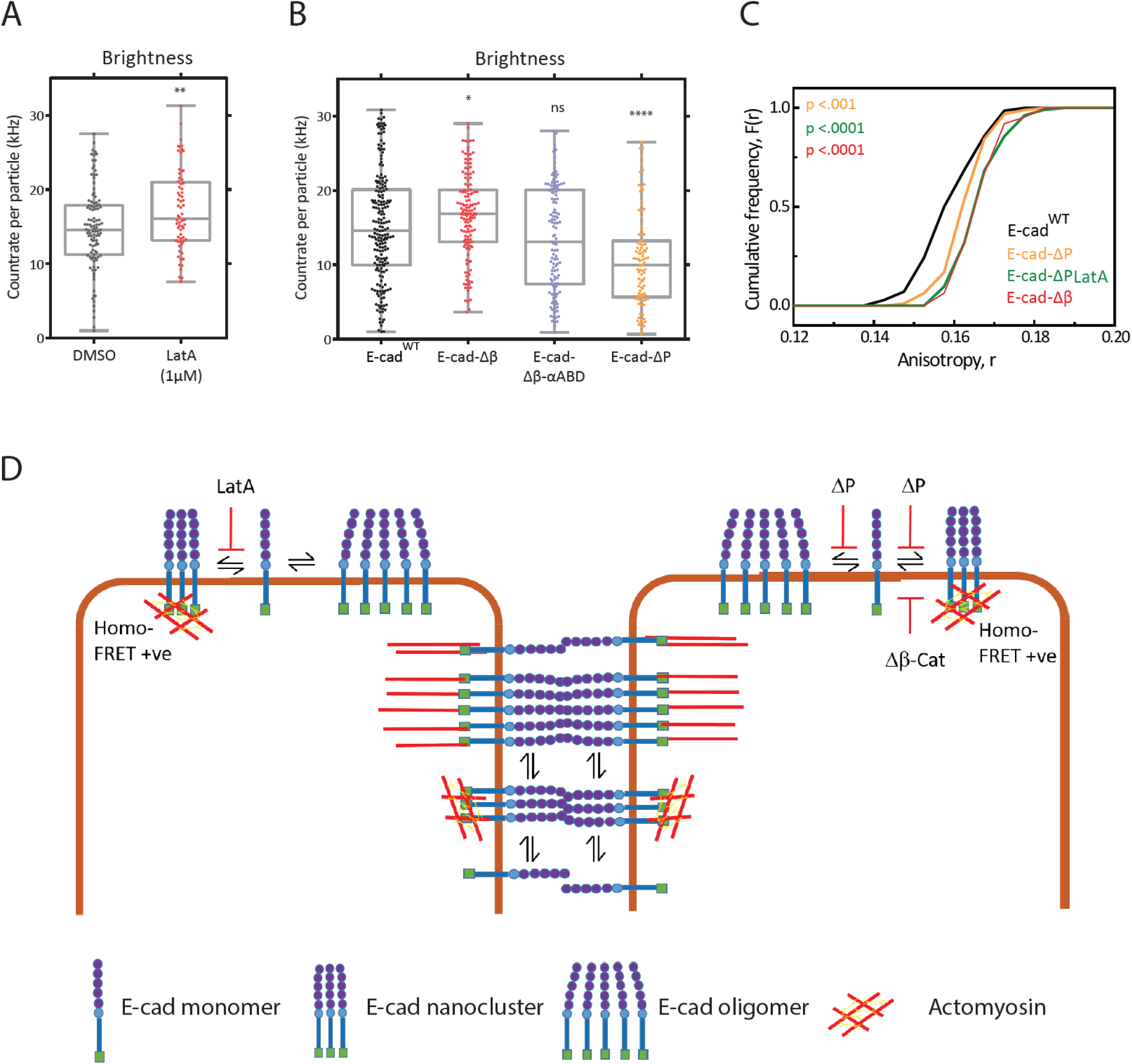
Analysis of E-cadherin oligomerization. (A) Brightness distribution of E-cad^WT^-GFP oligomers in cells treated with “DMSO” or “LatA (1µM)”. Other parameters estimated through related FCS analysis are detailed in Figure S6. (B) Brightness distribution of oligomers of various GFP tagged constructs of E-cadherin in cells expressing these constructs using UAS-GAL4 system. Other parameters estimated through related FCS analysis are detailed in Figure S7. (C) Cumulative frequency distributions of anisotropy values of lamellar E-cad in E-cad^WT^, E-cad-ΔP and E-cad-Δβ expressing hemocytes. (D) Schematic showing formation of *cis*-oligomers and actomyosin-sensitive nanoscale clusters in different conditions. p values are calculated using Mann-Whitney U test. ns, p>0.05; *, p<0.05; **, p<0.01; ***, p<0.001; ****, p<0.0001

### E-cadherin molecules exhibits a functional hierarchical organization

Having described the organization of extra-junctional E-cad molecules as consisting of at least two different states, namely, diffusive oligomers and relatively immobile acto-myosin dependent nanoclusters, we asked if this organization is hierarchical. There are two possibilities; i) a simple model to describe such a hierarchical organization would have the oligomers form spontaneously due to protein-protein interactions in *cis*, and these in turn promote nanoscale clusters, or ii) actively assembled nanoclusters could promote the formation of oligomers.

To test and distinguish between these alternatives we asked, how the nanoclustering status of E-cad-ΔP construct (which cannot form oligomers) compares with E-cad^WT^. Emission anisotropy measurements of E-cad-ΔP shows reduced nanoclustering as compared to E-cad^WT^ **(Figure 7C)**. Previously, we have established that the nanoclustering of E-cad requires an interaction with F-actin. In the case of E-cad-ΔP, the cytoplasmic domain of E-cad is intact; indicating that the nanoclustering is partially lost despite the potential for interaction with F-actin. Perturbation of actin interactions by treatment with LatA further reduces nanoclustering of E-cad-ΔP **(Figure 7C)** supporting the claim that E-cad-ΔP interacts with actin. Consistent with the first model, E-cad-ΔP also makes smaller diffusive oligomers as compared to E-cad^WT^ **(Figure 7B)**. This supports the hypothesis that *cis* interaction of E-cad is necessary for forming diffusive oligomers and for efficient acto-myosin-dependent E-cad nanoclustering.

E-cad makes larger diffusible oligomers when disconnected from actomyosin meshwork, as observed upon treatment of E-cad^WT^ expressing cells with LatA or for E-cad-Δβ expressing hemocytes **(Figure 7A, 7B)**. But in this state, the increased oligomeric nature (detected by FCS) does not result in a homo-FRET read out in the lamellar membrane **(Figure 4G)**. This suggests that in the absence of acto-myosin driven clustering at the cytoplasmic leaflet, E-cad forms a distinct type of large diffusible oligomeric species that are not sufficiently packed to elicit FRET between the GFP moieties. These oligomers are promoted by the *cis-*interactions of the ectodomain, revealing the presence of hitherto underappreciated types of E-cad species in the membrane. The dynamic nature of these *cis*-oligomers and the tendency of actin coupling to limit their growth also reveal an unanticipated function of actin in regulating the dynamic hierarchical organization of E-cadherin in free membrane and at cell junctions.

## DISCUSSION

In our study in hemocytes and the embryonic epithelium, utilizing fluorescence emission anisotropy and homo-FRET imaging, as well as FCS, we identify different types of higher-level organization of E-cad at the free membrane as well as at cell-cell junctions. We find that there are, 1) diffusive *cis*-oligomers in the lamella, 2) acto-myosin based *cis*-nanoclusters both at cell-cell junctions and in the contact free (lamellar or apical) membrane, and 3) *trans-*paired E-cad at cell-cell junctions.

We first consider the arrangement of the E-cad molecules at contact free surfaces, which are the lamellar surface in hemocytes and the apical surface in embryos. A consistent model that reconciles all our observations would have spontaneously interacting monomers form *cis*-oligomers to, in turn, facilitate actomyosin-sensitive nanoscale clusters (see Model in **Figure 7D**).

Evidence for this scheme wherein E-cad exists as two distinct species in the free surface comes from several observations. First, E-cad-Δβ is capable of forming large diffusive oligomers consisting of several (>4) E-cad monomers (**Figure 7B**), despite its inability to associate with cytoplasmic adaptor complexes. This interaction appears to be mediated by the ectodomain, since when the proximal domain of the ectodomain is perturbed, as in E-cad-ΔP mutant, the size of the diffusive species is further reduced (**Figure 7B)**. Second, the oligomers formed by E-cad-Δβ do not generate a measurable homo-FRET signal in the free surface (**Figure 4C**), indicating that the molecular arrangement of the cytoplasmic GFP/YFP tags are not within the FRET-range (Forster’s Radius ∼ 4-5 nm) in the diffusive oligomers. Third, engagement with the cytoplasmic acto-myosin machinery brings the cytoplasmic regions together creating nanoclusters of E-cad capable of exhibiting homo-FRET between the cytoplasmic fluorescent tags. Finally, perturbing the ectodomain interactions in the E-cad-ΔP mutant reduces the extent of actomyosin-based nanoclusters, and the size of the diffusive oligomers (**Figure 7C**). These observations suggest a hierarchical assembly mechanism for generating the acto-myosin based nanoclusters, wherein spontaneous *cis-*interactions of the E-cad protein facilitate the active nanoclustering step.

E-cad nanocluster organization requires coupling to an intact cortical acto-myosin network via the adaptor proteins, β-catenin, and the actin-binding α-catenin as well as the mechano-*trans*ducer vinculin [26]. The myosin regulator ROCK and the actin nucleator formin are also involved in this type of clustering [21, 43]. β-catenin binding mutant of E-cad, E-cad-Δβ, fails to form nanoclusters, and addition of the α-catenin ABD domain restore this clustering (**Figure 4C, 4D**). This confirms that in the context of nanoclustering, β-catenin is required as a link for E-cad to recruit α-catenin thereby connecting the protein to the actin network, also in accordance with the observations from other groups [40].

At cell-cell junctions in the hemocytes, E-cad is present in *trans*-paired assemblies, along with actomyosin-based nanoclusters. E-cad-Δβ also accumulates at the cell-cell junction in a manner that is capable of exhibiting measurable homo-FRET. However, here the nature and organization of the E-cad-Δβ at the cell-cell junction is very different from E-cad^WT^. Instead of having a population of nanoclusters and *trans*-paired species similar to E-cad^WT^, E-cad-Δβ exhibits *trans*-interactions and patches of densely packed proteins at the junction (**Figure 6H**). Evidence for *trans*-pairing comes from our experiments comparing the emission anisotropy of E-cad and E-cad-Δβ. E-cad-Δβ is more polarized at the junction **(Figure 6B)** compared to its counterpart in the lamella **(Figure S2G)**.

By contrast, E-cad-ΔP does not accumulate significantly at the junctions, and E-cad-ΔP is much less polarized than E-cad^WT^ **(Figure 2H; Figure S2H)**, indicating much weaker *trans-*interactions. Perturbation of actin reduces the extent of nanoclusters at the E-cad-ΔP junctions **(Figure S2H**), and E-cad-ΔP at the junction behaves somewhat similar to E-cad-ΔP lamella (**Figure S2I**). The formation of small diffusible oligomers and lack of junctional accumulation of protein at the junction in E-cad-ΔP indicates that *cis*-oligomerization of E-cad may be an important step in generating *trans* interactions at the junction. Alternatively, it is also possible that an intact proximal domain is required for strengthening *trans-*interactions.

The deletion of β-catenin binding domain (E-cad-Δβ) had no influence on the appearance and level of the mutant protein on the hemocyte membrane, but it affects its nanoclustering as well as its cell-cell interacting ability **(Figure 6B, 6C)**. Interestingly the cell adhesive strength of E-cad was affected when the interaction with adaptor components is disturbed since E-cad-Δβ or E-cad-Δβ-αABD expressing cells both had less cell aggregates compared to E-cad^WT^ expressing cells **(Figure 6C, 6F)**. Although E-cad-Δβ exhibits *trans* interactions, it lacks association with E-cad complex and F-actin. It is known that there is a force dependent strengthening of E-cad-E-cad interactions via α-catenin coupling to F-actin [6,13,48,49] and *trans* interactions of E-cad are likely to be strengthened by transmitting the tensile forces generated by association with the actomyosin cortex in adjacent cells [43].

The interaction with the actin cytoskeleton is restored in the E-cad-Δβ-αABD isoform since it exhibits extensive depolarization of its emission anisotropy at the lamella **(Figure 4D)**. However, this variant does not support robust cell-cell interactions **(Figure 6F)** exhibiting markedly enhanced nanoclustering propensity at the junction **(Figure 6E)**. These observations show that strong cell-cell interactions not only require *trans*-pairing of the *cis*-oligomers, but also require tunable interaction with cytoplasmic actin machinery necessary for junctional strengthening. These observations strongly suggest that the strength of intercellular adhesion does not simply reflect the strength of intra and intercellular molecular interactions, but their dynamic remodeling, consistent with the view that energy dissipation and rheology at adhesive interfaces is a critical element of adhesion [50]. In this context, it is likely that an actomyosin architecture strengthens *trans*-interactions and may act normal to the junction and exert tensile stress, e.g. the medial apical contractile actomyosin network in *Drosophila* epithelial cells, and in other organisms [43]. Moreover, the actomyosin nanoclustering machinery which acts naturally parallel to the junction, may assists in the disassembly of the *trans*-paired E-cad, for instance by exerting shear stress [43].

Our study in hemocytes suggests that actomyosin machinery helps in the nanoscale organization of junctional and extra-junctional E-cad, and is mirrored by similar observations in the embryonic epithelium, where both junctional and extra-junctional endo-E-cad::GFP appear to form nanoclusters that exhibit depolarized fluorescence emission anisotropy. Despite the usefulness of the hemocyte system in teasing out the hierarchical nature of E-cad structures at the nanoscale at the free surface of the hemocyte, the junctions in hemocytes compared to those in other epithelial cells is different. For example, unlike the embryonic epithelium, the cell-cell junctions do not seem to be under tension in the hemocytes-nor are they subject to active feedback. The junctions appear in a zig-zag pattern **(Figure 2C** [51]) which clearly indicate that they are not under tension and mechanisms that are responsible for junctional remodeling may not be operating in the hemocytes. E-cadherin based adhesion in epithelial cells is known to recruit the regulators actomyosin activity [52], which in turn regulate F-actin polymerization and Myosin-II activation to strengthen cell-cell adhesion. This might not be the case in hemocytes, since they do not express E-cad endogenously, and may not have the full-fledged machinery for self-regulation. Further exploration of the source of actomyosin regulation (junctional or extra-junctional) could point towards potential sources of E-cadherin dysregulation in epithelial cancers.

In conclusion, our characterization of diffusible and dynamic oligomers (loosely packed because they do not exhibit Homo-FRET) whose growth is restricted by actin coupling, as distinct from the tightly packed (Homo-FRET-competent) acto-myosin driven nanoclusters, reveal a complex regulatory role for the actin machinery on the nanoscale organization of E-cad. Actin coupling to loosely packed oligomers effectively reduces their size but also endows them with remodeling capacity which, we propose, is a prerequisite for their evolution to more stable actin dependent nanoclusters, and, eventually their *trans*-paired association. This may endow cells with the capacity to actively tune *trans* pair assembly by adjusting to the dynamically changing mechanical/contractile environment of a cell.

## ACKNOWLEDGMENTS

We thank H. Oda and P. Rørth for the gift of flies and plasmid constructs; Bloomington fly facility and Vienna Drosophila Resource Center for providing transgenic flies; Thomas Van Zanten for the help on FCS; Lokavya Kurup and Screening facility for cell aggregate imaging; D. Trivedi and NCBS fly facility for microinjection system and the Central Imaging and Flow Cytometry Facility at NCBS. We thank Mayor lab members for their valuable comments on the manuscript. R.C. acknowledges CSIR-UGC JRF fellowship from CSIR (Government of India). G.K. acknowledges bridging post-doctoral fellowship from NCBS. This project benefited support from the CNRS (Laboratoire International Associé, (SysTiM) between IBDM and NCBS) and Aix-Marseille Université, A∗MIDEX, ANR-11-IDEX-0001–02, funded by the Investissements d’Avenir French government program (BioTIM) and JC Bose Fellowship from DST (Government of India), Margadarshi fellowship (IA/M/15/1/502018) to S. M.

## AUTHOR CONTRIBUTIONS

R.C., T.L. and S.M. conceived the project. R.C. did EA and homo-FRET experiments and related data analysis. G.K. did FRAP and FCS experiments and related data analysis. J.-M.P. made the constructs and generated transgenic flies for this study. R.C., G.K. and S.M wrote the manuscript with comments from all the authors.

## STAR * METHODS

### KEY RESOURCES TABLE

| REAGENT or RESOURCE | SOURCE | IDENTIFIER |
| --- | --- | --- |
| **Antibodies** |  |  |
| Rat monoclonal anti-E-cadherin | DSHB | Cat#DCAD2,<br>RRID:AB_528120 |
| Mouse monoclonal anti-β-catenin | DSHB | Cat#N27A1,<br>RRID:AB_528089 |
| Rat monoclonal anti-α-catenin | DSHB | Cat#DCAT-1,<br>RRID:AB_532377 |
| Alexa Fluor 568 Phalloidin | Thermo Fisher Scientific | Cat#A12380 |
| Rabbit monoclonal anti-zipper | Lecuit Lab | NA |
| **Chemicals, Peptides, and Recombinant Proteins** |  |  |
| Ethyleneglycol-bis(2-aminoethylether)-N,N,N',N'-tetraacetic acid (EGTA) | Sigma Aldrich | Cat#E0396 |
| ROCK Inhibitor, H 1152 dihydrochloride | Tocris | Cat#2414 |
| Actin inhibitor, Latrunculin A | Thermo Fisher Scientific | Cat#L12370 |
| **Experimental Models: Organisms/Strains** |  |  |
| *D.melanogaster*: endoE-cad::GFP, sqh Sqh::mCherry | Sqh-mCherry gift from Adam C Martin | [53] |
| *D.melanogaster*: endoE-cadΔP::GFP/CyO | Lecuit Laboratory | N/A |
| *D.melanogaster*: VsV-G::GFP | Lecuit Laboratory | N/A |
| *D.melanogaster*: UAS-E-cad::GFP | Gift from H.Oda | [34] |
| *D.melanogaster*: Col-GAL4 | Gift from C. Dearolf | [54] |
| *D.melanogaster*: UAS-Ecad$\Delta\beta$cat::GFP | Lecuit laboratory | N/A |
| *D.melanogaster*: UAS-Ecad$\Delta\beta$cat-αcatABD::GFP | Lecuit laboratory | N/A |
| *D.melanogaster*: UAS-E-cad::YFP | Lecuit laboratory | N/A |
| *D.melanogaster*: UAS-Ecad$\Delta\beta$cat::YFP | Lecuit laboratory | N/A |
| *D.melanogaster*: <u>w<sup>1118</sup></u>; <u>P{GD1695}v7916</u> | Vienna Drosophila Resource Center | VDRC:v7916 |
| *D.melanogaster*: <u>w<sup>1118</sup></u>; <u>P{GD1566}v7819</u> | Vienna Drosophila Resource Center | VDRC:v7819 |
| *D.melanogaster*: UAS-EcadΔP | Gift from H.Oda | [36] |
| **Software and Algorithms** |  |  |
| MATLAAB, Image processing toolbox | Mathworks Inc | https://www.mathworks.com/products/matlab.html |
| Image J | N/A | https://imagej.nih.gov/ij/index.html |
| Metamorph 7.0 software | Molecular Devices Corporation, CA, USA | https://www.moleculardevices.com/products/cellular-imaging-systems/acquisition-and-analysis-software/metamorph-microscopy |
| GraphPad Prism | GraphPad | https://www.graphpad.com/scientific-software/prism/ |
| OriginPro 8 | Origin Lab | https://www.originlab.com/ |

### CONTACT FOR REAGENT AND RESOURCE SHARING

Further information and requests for resources and reagents should be addressed to and will be fulfilled by the Lead Contact, Satyajit Mayor.

### EXPERIMENTAL MODEL AND SUBJECT DETAILS

#### Hemocyte isolation and culturing

Hemocytes were cultured in Schneider’s complete medium (SCM) containing Schneider’s insect medium (Gibco-BRL, Gaithersberg, MD) supplemented with 10% non-heat inactivated serum (Gibco-BRL) and 1µg/ml bovine pancreatic insulin (Sigma-Aldrich). SCM was aged overnight at 4°C prior to use. Hemocytes were dissected from the third instar larvae as previously described (Sriram et al., 2003) in 150µl of SCM and were incubated in a BOD incubator at 23°C. Hemocytes were imaged 2.5-hour post-dissection.

#### Embryo Preparation

*Drosophila melanogaster* flies were maintained at 22^0^C in glass vials containing standard fly media. For embryo collection flies were maintained in cages with apple juice agar plates, supplemented with yeast paste. Embryos collected on these plates are washed with water and then treated with bleach to remove chorions. Early gastrulation embryos were aligned on the coverslips and covered with Halocarbon oil 200 for fluorescence imaging.

### METHOD DETAILS

#### Transgenic lines and genetics

*yellow white (y w)* flies were used as a control for immunostaining. UAS-Ecad::GFP and UAS-SHGDP#10 were gift from H. Oda. UAS-E-cad-Δβ::GFP, UAS-E-cad-Δβ-α-catABD::GFP, UAS-E-cad::YFP and UAS-Ecad-Δβ::YFP lines were generated in Lecuit lab. Collagen-GAL4 (gift from C. Dearolf) was used to drive expression of UAS constructs in hemocytes. endo-E-cad::GFP is a knock in line of E-cadherin at the locus (Huang et al, 2009). Myo-RLC::mcherry (gift from A. Martin) is also imaged along with E-cad::GFP in embryo for drug perturbations. endo-E-cad-ΔP/CyO flies was generated in Lecuit lab. Myosin-II RNAi lines, MRLC RNAi and MHC RNAi, were obtained from VDRC. Ubi-VsVg::GFP line (Thomas Lecuit) was used as control for anisotropy measurements in embryo.

#### Drug injection

Embryos were prepared as described before. For drug injections, embryos were kept in box containing Drierite and injected with either drugs or similar volumes of their solvents (RNase free water or DMSO), at the end of cellularisation or early gastrulation. For ROCK inhibition, embryos were injected with 20mM H1152 (Tocris) and imaged immediately. For actin perturbations, embryos were injected with 5mM Latrunculin A (Invitrogen) and imaged immediately. For anisotropy measurements, 15-20 Z-sections separated by 0.25µm were acquired from the most apical membrane of the embryo.

#### Actomyosin perturbations in hemocytes

Actin perturbations were carried out using 20µM Latrunculin A (Invitrogen) for 45 minutes. Formin perturbation was carried out using 25μM SMI-FH2 (Sigma Aldrich) for 1 hour. ROCK perturbations were carried out using 10µM H1152 (Sigma) for 1 hour. Imaging was carried out in the presence of the drug. All drug treatments were carried out in M1 buffer (150mM NaCl, 5mM KCl, 1mM CaCl2, 1mM MgCl2 and 20mM HEPES – pH 6.9) containing 1mg/ml BSA and 2mg/ml glucose at 23^0^C BOD incubator for the indicated time periods.

#### Immunofluorescence imaging/microscopy

Hemocytes were fixed with 2.5% paraformaldehyde for 20 minutes at room temperature and permeabilized with 0.37% Igepal for 13 minutes at room temperature for whole cell protein labeling. Cells were blocked using M1 containing 2mg/ml BSA and 2mg/ml glucose (blocking buffer) for 30 minutes and labeled with rat anti-E-cadherin (1:10 DCAD2, DSHB), mouse anti-beta catenin (1:10 N27A1, DSHB), rabbit anti-alpha catenin (1:10 DCAT2, DSHB) or rabbit-anti zipper (1:150, Gift from Thomas lab) for 1 hour and then labeled with corresponding fluorescent secondary antibody. For actin labeling cells were labeled with Alexa 568 labeled phalloidin (Molecular probes). Immunostained samples were imaged using a wide field microscope (Nikon) using a Plan Apochromat 60X/1.4 NA oil immersion objective.

#### Fluorescence Emission Anisotropy measurements and image analysis

Fluorescent samples were excited by plane polarized light and the emission was split into parallel (I_pa_) and perpendicular (I_pe_) components with the help of a polarizer fitted in the path and collected simultaneously by two EMCCD cameras. Fluorescence anisotropy was calculated from the intensities collected in the parallel and perpendicular cameras using the formula

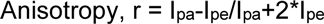

Anisotropy measurements of hemocytes were carried out in TIRF microscope setup on Nikon TE2000 body, having polarized 488 laser, using a 100X objective with NA of 1.49 [28]. Embryo measurements were carried out in confocal Spinning disk microscope equipped with a Yokogawa CSU-22 unit using 100X objective with NA of 1.4. Andor laser combiners emitting 488 and 561 nm were used to image embryos and the intensities were collected using Andor ixon+897 EMCCD cameras[28]. Photobleaching experiments were done in TIRF microscope using stream acquisition with an interval of 50-100ms between frames.

The parallel and perpendicular fluorescent images were background subtracted using M1 buffer image taken at the same imaging conditions. Both images were then aligned using Matlab code in MATLAB (Mathworks, USA) and G-factor corrected. Hemocyte lamellar anisotropy measurements were taken from multiple 20×20 pixel (1.7*1.7 µm) regions selected on the flat lamellopodium of the hemocyte using Metamorph^TM^ software. Hemocyte junctional measurements were taken by dividing the junctions into multiple small regions. The intensities of parallel and perpendicular images were extracted and anisotropy values were calculated using the formula described above. Anisotropy values of similar total intensity range are plotted as cumulative frequency distributions in Origin software. Anisotropy maps of the cells were also generated from the aligned images using a code written in Matlab.

Embryo images were also processed through the same steps before anisotropy measurements. Apical anisotropy measurements were taken from 5×5 pixel (0.68 × 0.68 µm) regions from the apical most membrane of the embryo. Junctional punctae measurements were taken from junctional membrane plane, which is approximately 1µm from the apical most membrane. Punctae measurements were taken from around 5-6 Z-planes, separated by 0.25µm, of the junctional membrane. For junctional punctae analysis, images were again subtracted using a 50-pixel radius median filter. Embryos depending on the stage or its generation (F0 vs F2) were having different medial intensities and were affecting the anisotropy values. Since different embryos were having different medial intensities, this step was introduced to reduce the effect of medial pool on the anisotropy measurements.

For photobleaching analysis on hemocyte lamella and junction, 20×20-pixel (1.7*1.7 µm) regions selected on the flat lamellopodium for lamellar measurements and regions were drawn on the junctions for junctional measurements. Intensities and anisotropy values of all frames were calculated using the formula described above. The intensities in the other frames were normalized to the intensity in the first frame of the ROI. Scatter plot of average anisotropy values of different ROIs from different cells against binned normalized intensity ratios was plotted.

#### Fluorescence Recovery After Photobleaching (FRAP)

We used Zeiss LSM 780 Confocor 3 System for the FRAP experiments using a 40X 1.2 NA UV-VIS-IR C Achromat water-immersion objective. The laser power at the back-focal plane of the objective was kept constant at 6uWfor the 488nm line from Argon-laser. A circular ROI (diameter 3µm) was photobleached by switching the laser power to 100%. Another reference ROI of the same dimensions was drawn in the same cell to estimate imaging related photobleaching. A third ROI outside the cell was used to account for background. Images were acquired every 500ms, with 5 frames prior to photobleaching, and 120 frames after. The intensity traces in the bleach region were corrected for background and photobleaching, and were normalized using the averaged intensity in the pre-bleach region. Individual FRAP trace profiles were fitted to:

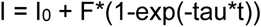

where, I is intensity at time-point t post-photobleaching. The bleaching is never 100% and the residual initial intensity is estimated through I_0_. The intensity recovers till the mobile fraction (F) is exchanged, at characteristic half-life ln2/tau. Here we are more interested in the immobile fraction which is estimated as 1-(F+I_0_).

#### Fluorescence correlation spectroscopy (FCS)

We used a point scanning confocal; Zeiss LSM 780 Confocor 3 System for the FCS measurements using a 40X 1.2 NA UV-VIS-IR C Achromat water-immersion objective. The back-focal plane of the objective was overfilled using 488nm line from an Argon-laser (at 6 uWatt, corresponding to about 5-10 kWatt/cm2) in order to create a diffraction-limited confocal volume that was calibrated on each day before the experiment by maximizing the count-rate per particle of Rhodamine 6G [55, 56]. Hemocytes were cultured and pretreated as describe above, depending on the experiment. Confocal spot was focused on the ventral membrane of the hemocytes in the lamellar region of the cell. The confocal spot was parked in the center of the field and region of interest in individual cell was moved there by moving the stage. The correct focal distance was determined each time as the z-distance where the initial estimate of counts per molecule were highest. Next, the emission photon stream was recorded with the same objective, descanned, through an aligned pinhole (32 µm), wavelength selected between 491-571 and detected on a gallium arsenide detector array.

Two intensity in time traces (*I(t)*) of 60 seconds each were recorded for each cell as 6 iterations of 10 seconds. Each 10 second trace was autocorrelated into an autocorrelation curve G(*τ*) using the Zeiss onboard autocorrelator which calculates the self-similarity through:

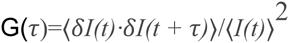

Here ⟨⟩ denotes the time-average, *δI(t)* = *I(t)* – ⟨*I(t)*⟩ and *τ* is called the timelag. The 10 second measurement is long enough to contain sufficient events and short enough to avoid each trace to be contaminated by events that do not arise from Ecad::GFP diffusing in-plane of the plasma membrane. 1ms-binned intensity traces were manually assessed for data quality. Traces that contained monotonous changes in intensity (most likely arising due to photobleaching and/or z-drift during the measurement) or contained short high-intense bursts (most likely arising from endosomes passage) were discarded from further analysis. If the first set of traces were useful upon visual inspection, another set of traces were recorded the same point in the cell. This way, each cell yielded either 12 or 24 traces of 10 seconds each, along with corresponding autocorrelations generated by the Confocor.

A potential distribution of fluctuation timescales was previously determined [46] for membranous proteins. This article indicated existence of 3 timescales as follows. First, a 20-120 us timescale corresponding to the triplet state of EGFP [29]; then second, a 0.5-5ms timescale corresponding to the intracellular/ luminal GFP that has also been observed previously in cells expressing EGFP tagged membrane protein [30, 57]; and third, a timescale of ranging 10-100ms corresponds to the lateral diffusion of membrane targeted GFP through the confocal spot.

We manually went through all 1ms binned intensity traces to discard the ones that show artifacts as mentioned above. For the traces that remain, we fitted their corresponding raw 10 s autocorrelation. The autocorrelation G(*τ*) was fitted to:

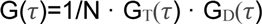

N reflects the number of moving particles in the confocal volume (including both 2D and 3D diffusing species) and G_T_(*τ*) is correlation function associated to blinking/triplet kinetics:

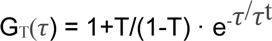

Where T is the fraction of molecules in the dark state and *τ*t corresponds to the lifetime of the dark state. G_D_(*τ*) is correlation function associated to diffusion, which in this case contains two diffusional timescales corresponding to either 2D or 3D diffusion as follows:

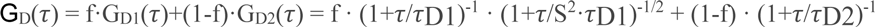

The fraction f corresponds to the intracellular 3D diffusing pool of EGFP that has a timescale of *τ*_D1_. S is the structure factor that accounts for timescales arising from the fact that the intracellular EGFP diffuses in a volume rather that a plane. Free fitting this parameter converges the value to about 0.2, consistent with earlier reports on cultured cells. In order to contain the number of free parameters we decided to fix this value to 0.2. The diffusion time associated with laterally diffusing Ecad::GFP is finally calculated with *τ*_D2_. When fitting the autocorrelation traces for GFP in solution or cytoplasm, the 2D component was excluded from the fit equation.

For estimating the brightness of 2D diffusing species of E-cad^WT^ or its variants, we first used the fraction f to estimate the number of diffusing entities in 2D and 3D as:

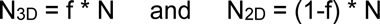

Then, solving following equation gives the brightness of the 2D diffusing component Q_2D_:

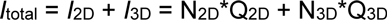

when, we plug in the brightness of cytosolic monomeric EGFP in place of Q_3D_, which was estimated in an independent experiment.

### QUANTIFICATION AND STATISTICAL ANALYSIS

Each hemocyte anisotropy experiment reported here has dataset from one experiment and was performed at least twice with similar results. For lamellar data quantification, a 20×20 pixel (1.7×1.7 µm) region of interest (ROI) based analysis was performed on 10-20 cells. For junctional data quantification, different area regions were drawn over 10-20 junctions in each experiment. Apical and junctional data quantification was done on 5×5 pixel (0.68 × 0.68 µm) region of interest drawn on embryos with same experimental conditions. Embryo experiments has been conducted over different days and the data was generated by pooling from different days embryos. N refers to the number of embryos taken for measurements.

Non-parametric Mann-Whitney test was performed to check the statistical significance between different experimental conditions and to calculate p values. p values are indicated in the figure panel itself. ns, p>0.05; *, p<0.05; **, p<0.01; ***, p<0.001; ****, p<0.0001

---

**Figure S1:**
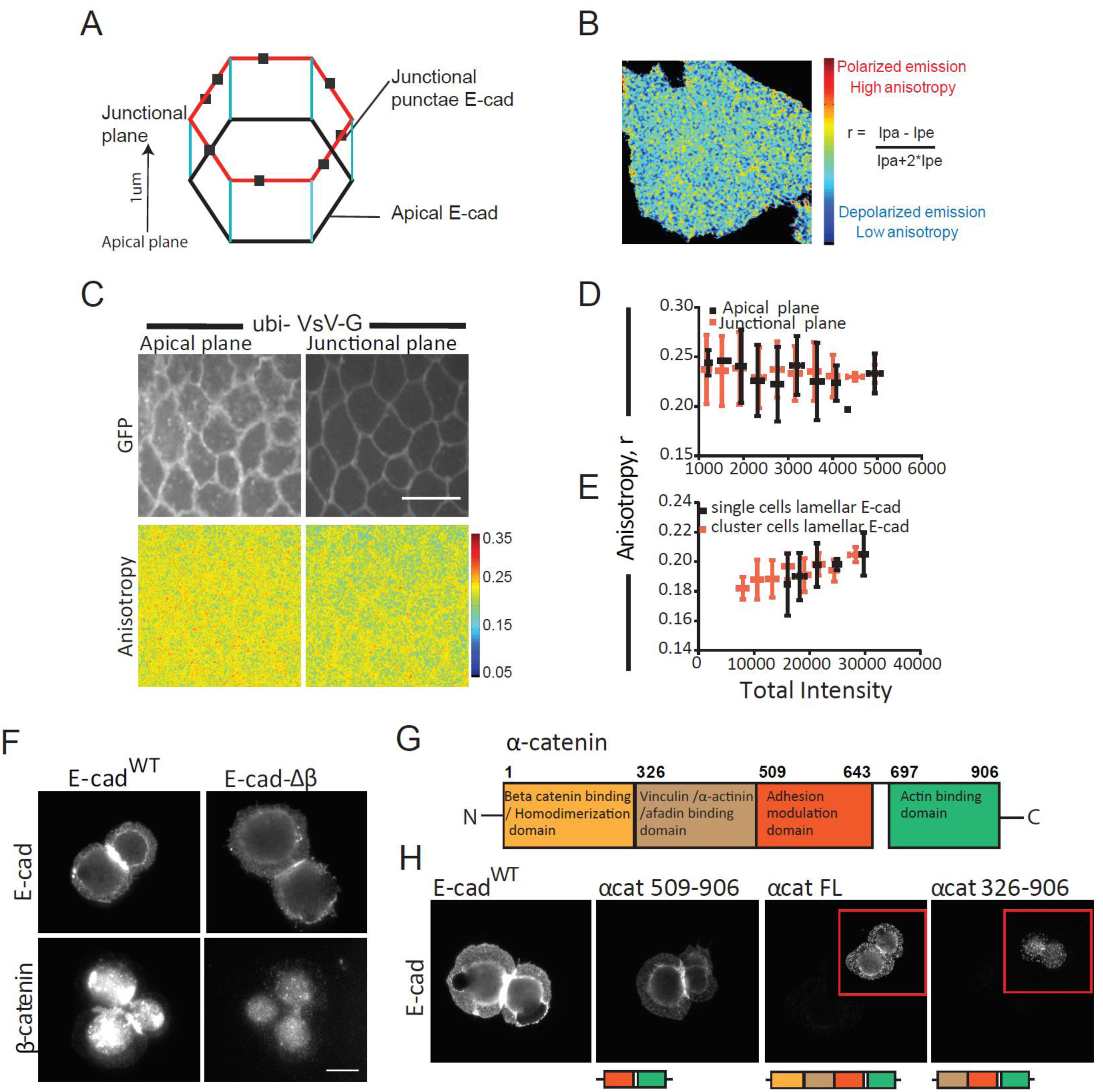
Transmembrane protein, VsV-G organizes in the embryo, and E-cad mutant proteins are functionally active in hemocytes. (A) Representation of epithelial cells of the embryo showing the planes at which apical and junctional anisotropy measurements are taken. (B) Schematic representing fluorescence anisotropy measurements. (C) Total intensity and anisotropy images of ubiquitous VsV-G::GFP in the apical and junctional plane of the early gastrulation stage of the embryo. Scale bars, 10μm. (D) Intensity versus anisotropy plots of ubi-VsV-G in the apical and junctional plane from similar stage embryos. (E) Cumulative frequency distributions of anisotropy values of lamellar E-cad from single cells and cluster cells. (F) E-cadherin and β-catenin localization in E-cad^WT^ and E-cad-Δβ expressing hemocytes as labeled with DCAD2 and N27A1 antibody. Scale bars, 10μm. (G) Schematic showing the key domains of α-catenin protein. (H) E-cad surface staining in different E-cad-Δβ-α-cat chimeric proteins expressing hemocytes as labeled with DCAD2 antibody. Images are equally scaled.

**Figure S2:**
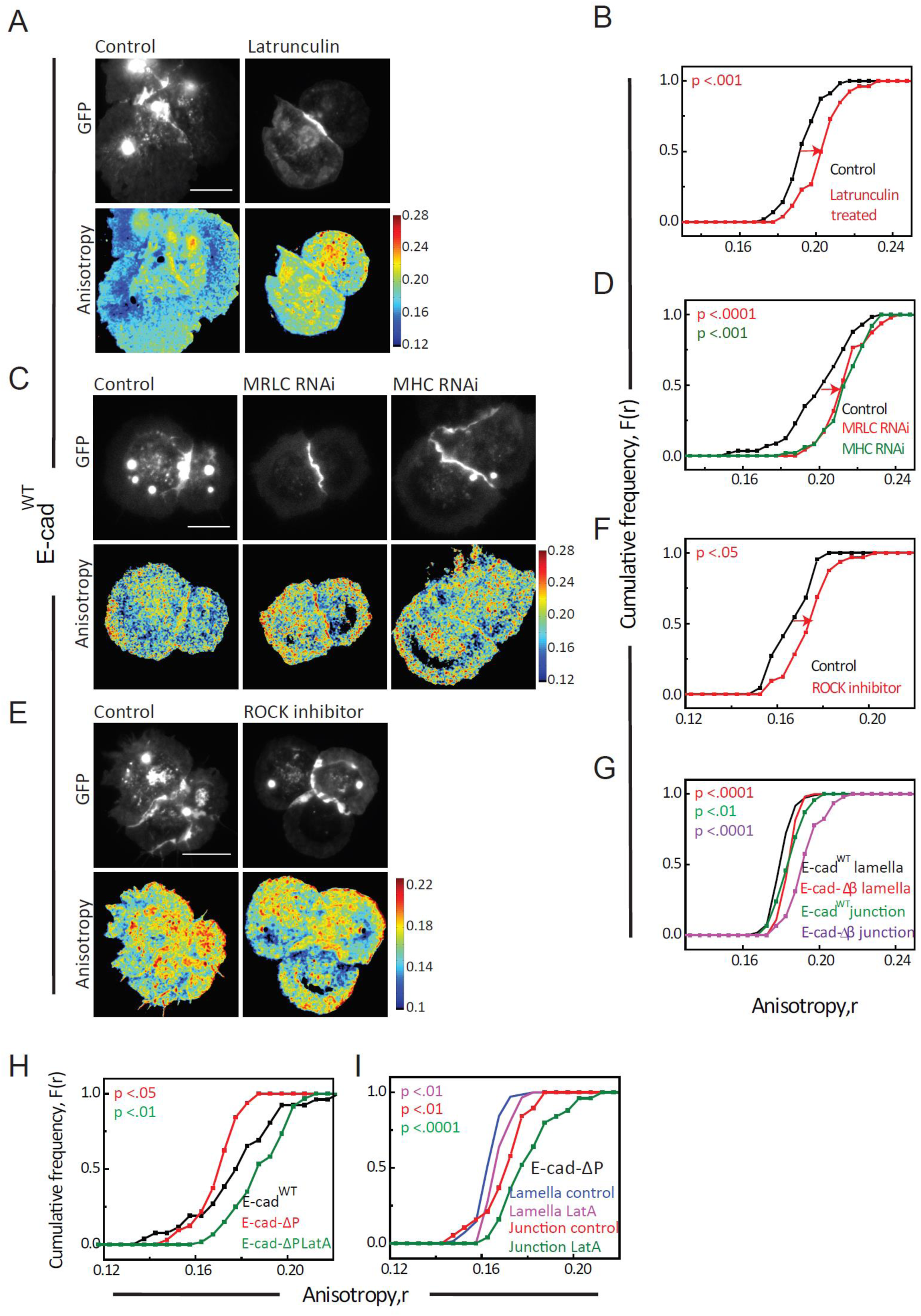
Acto-myosin dependent junctional organization of E-cad and its regulation. (A) Total intensity and anisotropy images of E-cad^WT^::GFP in Latrunculin A treated cells. Scale bars, 10μm. (B) Cumulative frequency distributions of anisotropy values of E-cad^WT^ in control and Latrunculin A treated cells. (C) Total intensity and anisotropy images of E-cad^WT^::GFP in control, MRLC RNAi and MHC RNAi expressing cells. Scale bars, 10μm. (D) Cumulative frequency distributions of anisotropy values of E-cad^WT^ in control, MRLC RNAi and MHC expressing cells. (E) Total intensity and anisotropy images of E-cad^WT^::GFP in control and ROCK inhibitor treated hemocytes. Scale bars, 10μm. (F) Cumulative frequency distributions of anisotropy values of E-cad^WT^ in control and ROCK inhibitor treated cells. (G) Cumulative frequency distributions of anisotropy values of E-cad^WT^::GFP and E-cad-Δβ::GFP at lamella and junction. (H) Cumulative frequency distributions of junctional anisotropy values of E-cad^WT^ control and E-cad-ΔP in control and Latrunculin A treated cells. (I) Cumulative frequency distributions of anisotropy values of E-cad-ΔP lamella and junction in control and Latrunculin A treated cells. p values are calculated using Mann-Whitney U test.

**Figure S3:**
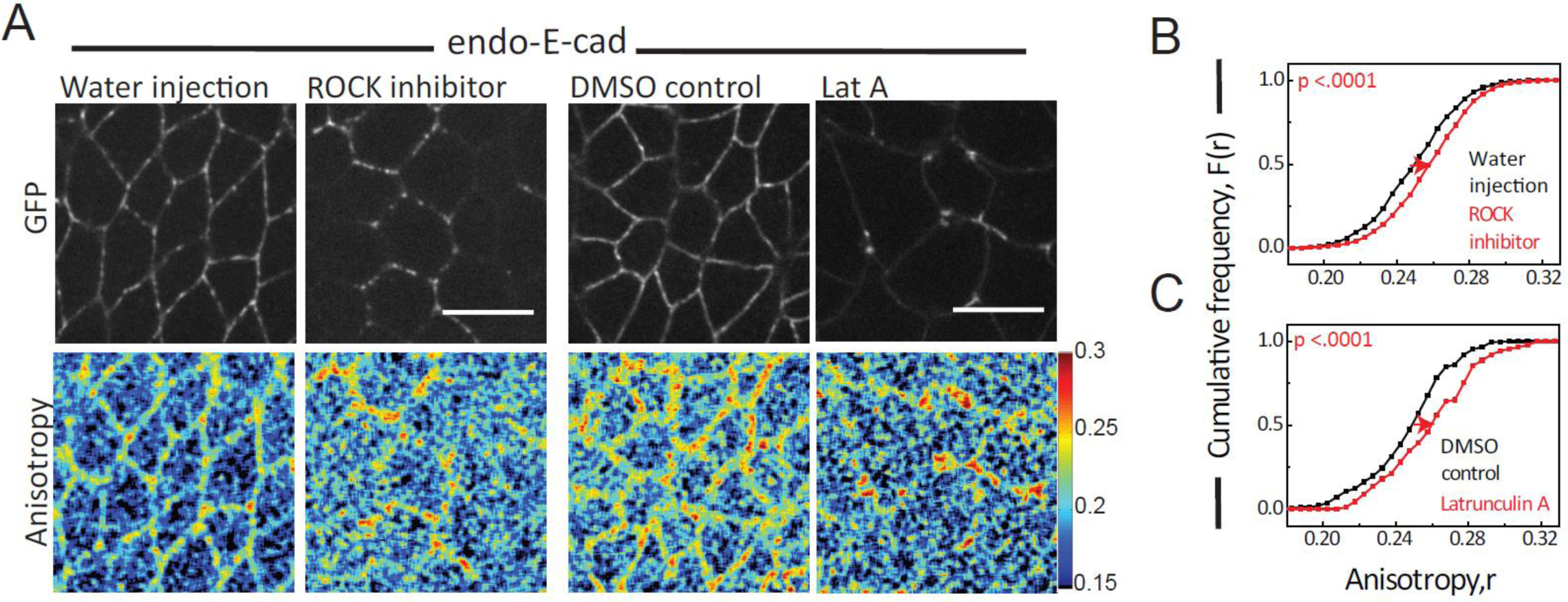
Nanoscale organization of *Drosophila* junctional E-cad is acto-myosin dependent. (A) Total intensity and anisotropy images of endo-E-cad in the junctional plane in the early gastrulation stage of the embryo, either control or micro injected with ROCK inhibitor or Latrunculin A. Scale bars, 10μm. (B) Cumulative frequency distributions of anisotropy values of junctional endo-E-cad in water injected (N=4) and ROCK inhibitor injected (N=6) embryos. (C) Cumulative frequency distributions of anisotropy values of junctional E-cad in DMSO injected (N=3) and Latrunculin injected (N=4) embryos. p values are calculated using Mann-Whitney U test.

**Figure S4:**
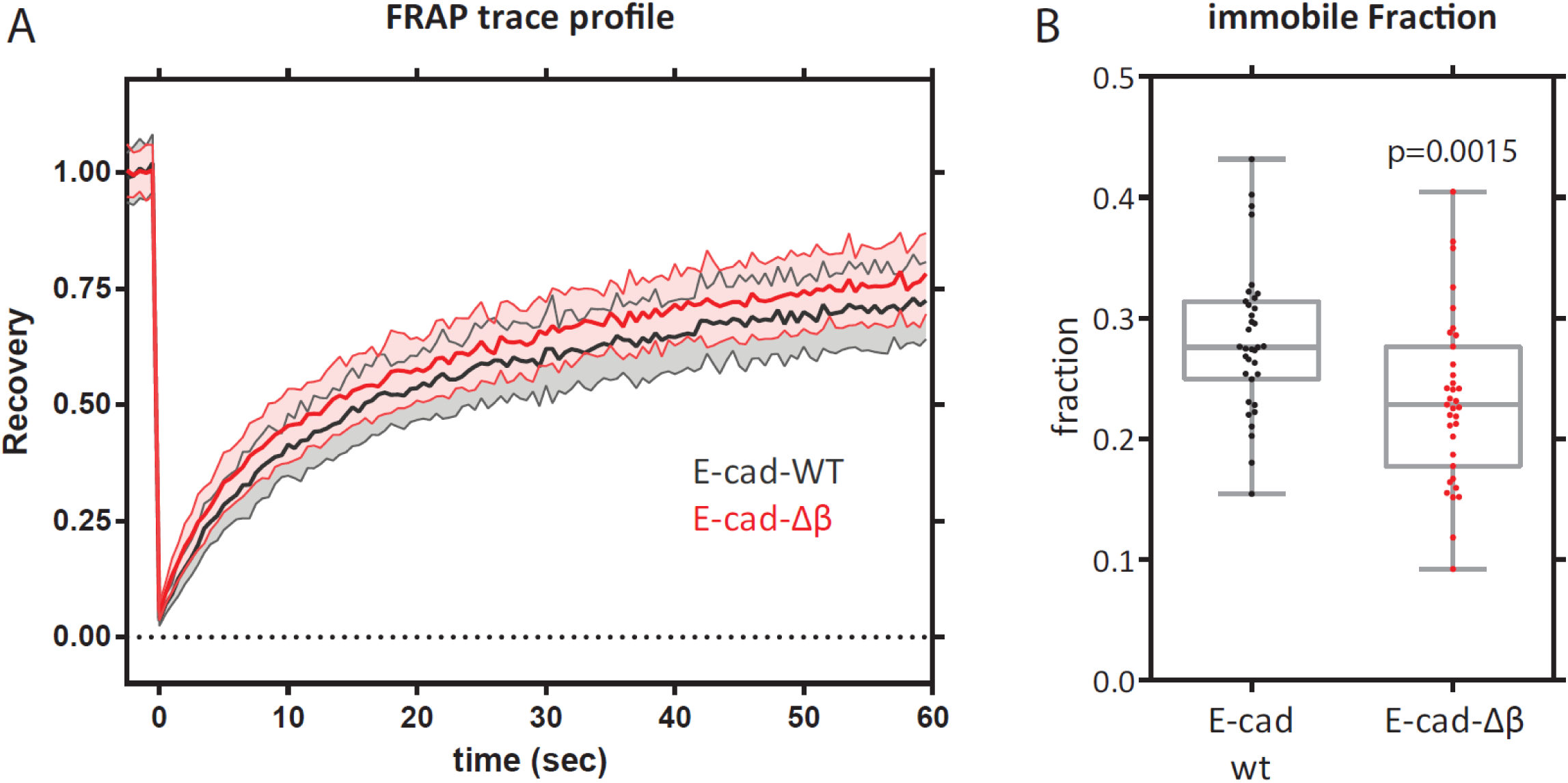
FRAP experiments in hemocytes. (A) FRAP trace profile showing the extent of fluorescence recovery as a function of time post bleaching in E-cad^WT^::GFP or E-cad-Δβ::GFP expressing cells. The bold line represents mean and shaded regions represent SD. (B) The estimated immobile fraction (see methods) for E-cad^WT^::GFP and E-cad-Δβ::GFP. Statistical significance estimated using Mann-Whitney U test.

**Figure S5:**
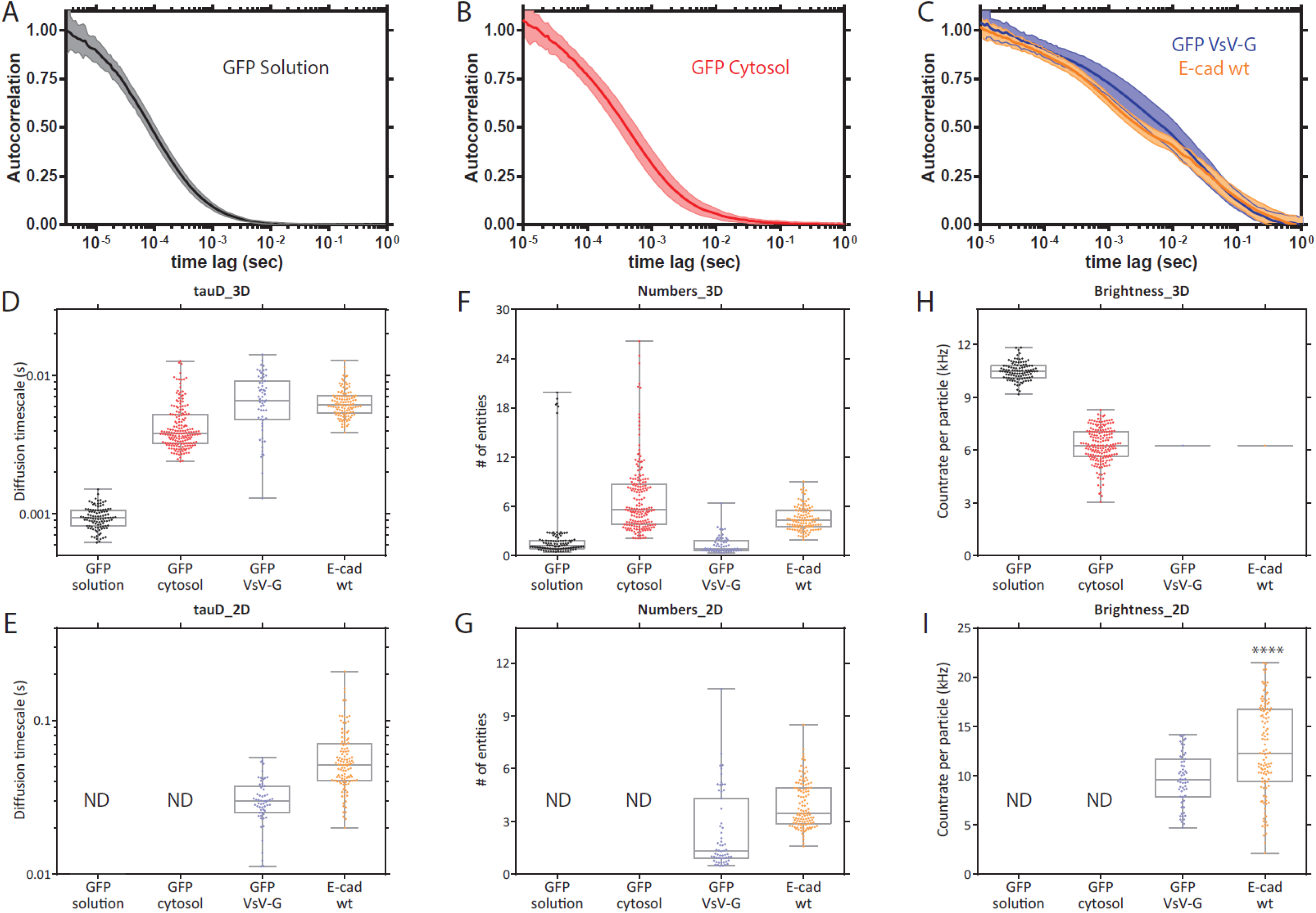
Supplementary figure for GFP intensity calibration FCS experiments. (A, B, and C) Autocorrelation traces for GFP in solution (A), in cytoplasm (B), and tethered to membrane via VsV-G or E-cad (C). The bold line represents mean and shaded regions represent SD. (D and E) Distribution of ‘diffusion time scales (tauD)’ for the 3D fraction (D) and the 2D fraction (E), estimated by fitting the respective individual autocorrelation traces. (F and G) Distribution of ‘number of diffusing species’ for the 3D fraction (F) and for the 2D fraction (G), estimated by fitting the respective individual autocorrelation traces. (H and I) Distribution of ‘brightness of diffusing species’ for the 3D fraction (H) and the 2D fraction (I), estimated by fitting the respective individual autocorrelation traces. The brightness estimates indicate that the cytoplasmic GFP has about half the brightness as compared to GFP dissolved in Phosphate Buffered Saline solution. Further, E-cad::GFP oligomers are significantly brighter than VSV-G::GFP (p<0.0001, Mann-Whitney test), and resemble a tetramer as they are about 1.25 fold brighter than VSV-G::GFP (which is a trimer). In E, G, and I, ‘ND’ indicates ‘no data’, as the GFP in solution or cytoplasm is not supposed to have a 2D diffusive component.

**Figure S6:**
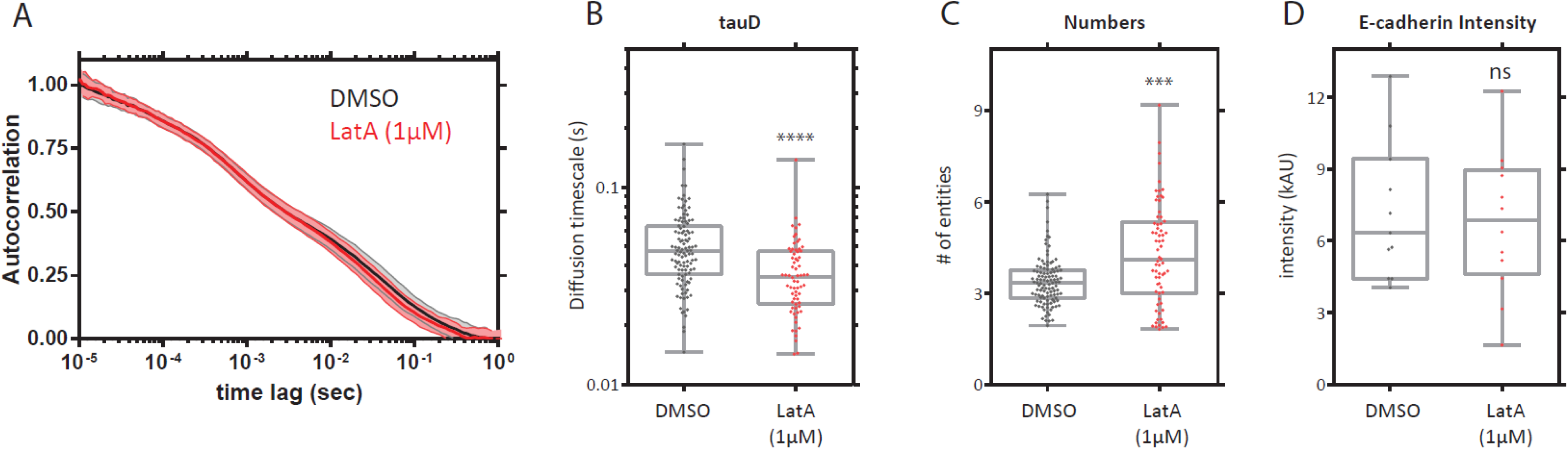
Supplementary figure for FCS experiments with LatA treatment. (A) Autocorrelation traces from E-cad^WT^ expressing cells treated with DMSO, as compared to treatments with 1µM LatA. The bold line represents mean and shaded regions represent SD. (B) Distribution of ‘diffusion time scales (tauD)’ for the 2D fraction, estimated by fitting the respective individual autocorrelation traces. (C) Distribution of ‘number of diffusing species’ for the 2D fraction, estimated by fitting the respective individual autocorrelation traces. (D) Distribution of GFP intensity at the FCS spot prior to FCS recordings (one data point per cell). p values indicate comparison with DMSO treatment and are calculated using Mann-Whitney U test. ns, p>0.05; *, p<0.05; **, p<0.01; ***, p<0.001; ****, p<0.0001

**Figure S7:**
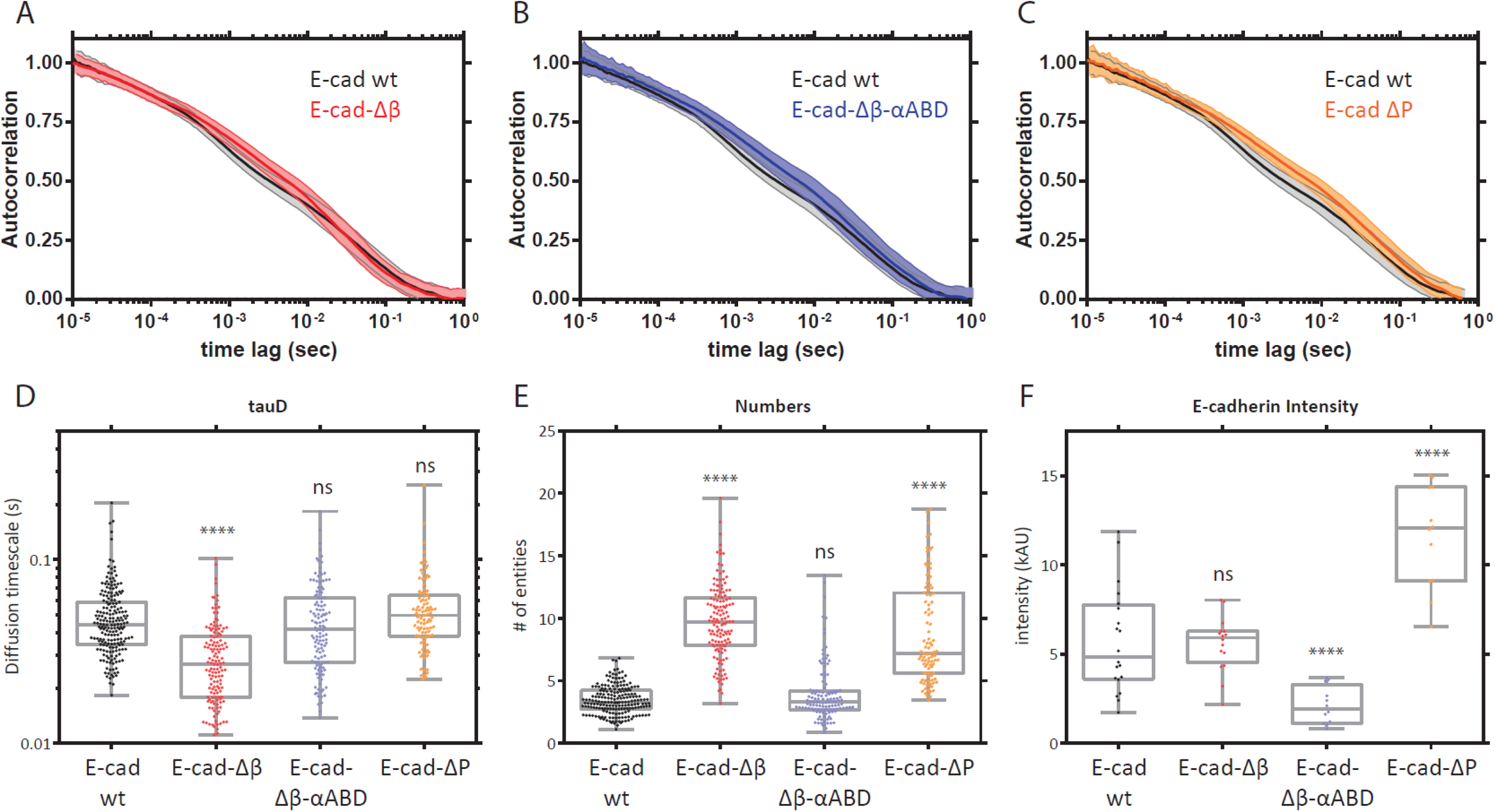
Supplementary figure for FCS experiments with E-cad deletion or chimeric constructs. (A, B, and C) Autocorrelation traces for E-cad^WT^ as compared to E-cad-Δβ (A), E-cad-Δβ-αABD (B), and E-cad-ΔP (C).The bold line represents mean and shaded regions represent SD. (D) Distribution of ‘diffusion time scales (tauD)’ for the 2D fraction, estimated by fitting the respective individual autocorrelation traces. (E) Distribution of ‘number of diffusing species’ for the 2D fraction, estimated by fitting the respective individual autocorrelation traces. (F) Distribution of GFP intensity at the FCS spot prior to FCS recordings (one data point per cell). p values indicate comparison with E-cad^WT^ and are calculated using Mann-Whitney U test. ns, p>0.05; *, p<0.05; **, p<0.01; ***, p<0.001; ****, p<0.0001

